# p53 supports healthy aging

**DOI:** 10.64898/2026.11.16.766897

**Authors:** Marcos G. Teneche, Karl N. Miller, Kelly Yichen Li, Kat Lande, Sainath Mamde, Christina Huan Shi, Adarsh Rajesh, Aaron P. Havas, Jessica Proulx, Rouven Arnold, Michael Alcaraz, Andrew Davis, Armin Gandhi, Laurence Haddadin, Xue Lei, Irene Huang, Gabriele Guarnaccia, Jake Castillo, Monica Sevilla, Hiroshi Tanaka, Cray Minor, Caroline Kumsta, Tatiana Moreno, Erik Hultenius, Carolina Cano Macip, Garrett Evensen, Shreeya Patel, Colin Kern, Elizabeth Smoot, Jacqueline Olness, Andrea Rodríguez-Agustín, Guillermina Garcia, Yu Xin Wang, Kevin M. Tharp, Alessandra Sacco, Rebecca A. Porritt, Quan Zhu, Allen Wang, Bing Ren, April Williams, Nathan Zemke, Kevin Y. Yip, Peter D. Adams

## Abstract

p53 is a master regulator of cellular stress responses that is also implicated in the aging process. Its role in aging is often considered an example of antagonistic pleiotropy (AP), where it acts beneficially as a tumor suppressor early in life, but detrimentally in late life, for example by driving accumulation of stress-associated, pro-inflammatory senescent cells. However, the causal role of p53 in aging is not well-defined, and the AP model has not been definitively tested through loss-of-function studies. Here we show, contrary to the prediction of AP, that p53 promotes healthy aging in mouse liver by suppressing excessive inflammation and maintaining metabolic function. Multimodal RNA, protein, epigenetic and spatial profiling at the single-cell level in aged mouse liver confirmed age-dependent increase in p53 pathway activation. Hepatocyte-specific knockout of p53 in livers of aged mice, but not of young mice, increased inflammation and immune infiltration. Mechanistically, p53 suppressed DNA replication stress and DNA damage, known precursors to inflammatory signaling. In aged hepatocytes, p53 also directly targeted and maintained expression of metabolic genes, and knockout of p53 sensitized aged mice to metabolic stress-induced liver damage. Complementing these loss-of-function data, pharmacological activation of p53 with the small-molecule MDM2 inhibitor HDM201 in aged mice reversed age-associated suppression of metabolic genes, decreased age-associated inflammation and delayed onset of frailty. These results identify p53 activity in aged tissue as protective rather than degenerative. Importantly, this protective function is pharmacologically targetable, potentially for the benefit of healthy aging in humans.

## Introduction

The tumor suppressor p53 has a well-established role as a “guardian of the genome”^1^, because of its central function in preserving genomic stability and preventing malignant transformation. Cellular stress activates p53, which in turn induces a transcriptional program including upregulation of the cyclin-dependent kinase inhibitor (CKI) p21 (CDKN1A), which enforces cell-cycle arrest^2–5^. Depending on the nature of the damage and stress response, p53 can drive apoptosis^6^ or entry into cell senescence^7,8^, both potent tumor suppressor mechanisms^9^. By coordinating these stress responses, p53 preserves genomic integrity and acts as a major barrier against malignant transformation. The importance of that barrier is underscored by *TP53* being the single most frequently mutated gene across human cancers^10^.

Beyond its canonical role in tumor suppression, p53 is recognized as a master regulator of organismal physiology, intersecting with most of the hallmarks of aging^11–13^. p53 target gene expression increases with age in non-diseased conditions across many tissues in mouse, rat, macaque and humans^14,15^. This activation is thought to reflect cumulative genotoxic, metabolic, and other stresses that accrue over the lifespan^11^. p53 is engaged by, and in turn modulates telomere attrition, mitochondrial function, autophagy and proteostasis, as well as nutrient sensing and metabolism^9,11^. It also influences the epigenome, stem cell renewal^16–18^, immune surveillance, and the clearance of damaged cells^19,20^. As an inducer of cellular senescence, p53 can also impact inflammatory programs, including the senescence-associated secretory phenotype (SASP), a proinflammatory secretory program of senescent cells^21–23^, and upregulation of secreted interferon-stimulated genes (ISGs) across most aged organs^15^. SASP and ISGs contribute to “inflamm-aging”, chronic inflammation of aged tissues that is thought to be a driver of age-associated disease^24,25^. These observations and diverse activities of p53 suggest that p53 is not only critical for preventing cancer but may also be a central determinant of the aging process itself.

The antagonistic pleiotropy (AP) theory^26^ proposes that aging is driven by pleiotropic gene action, whereby genes with beneficial functions early in life drive dysfunction late in life. Tumor suppressors are canonical examples of AP. For example, AP proposes that *TP53* prevents cancer in early life but may induce degenerative features of aging later in life, potentially through SASP and chronic inflammation^13,27^. Senescent cells accumulate with age^28^ and are increasingly considered drivers of age-associated tissue dysfunction^29–31^. However, although p53 induces senescence associated cell cycle arrest^32^, p53 inhibits the SASP^21,33,34^, through a mechanism involving p53-dependent DNA repair^34^.

Moreover, transgenic mouse models with increased p53 activity have not yielded consistent results regarding tumor and aging phenotypes^13,27,35–37^. Thus, the AP role for p53 is not universally supported. To better address this issue, the role of p53 in aging should be probed by a loss-of-function model; however, whole body p53 knockout (KO) mice are not informative for natural aging, because they develop cancer relatively early in life and are short-lived^38,39^. Here, we investigated the role of p53 in aging, using genetic KO and pharmacological approaches in aged mice. We show that, although p53 signaling is activated with age, KO of p53 in aged, but not in young, mouse liver drives inflammation and metabolic dysregulation. Using a complementary approach, activation of p53 with a small-molecule MDM2 inhibitor in aged mice delays phenotypes of aging. We propose that p53 activation with age is not detrimental; rather, it has beneficial functions that are pharmacologically targetable to promote healthy aging.

## Results

### p53 activity increases in aged hepatocytes

p53 activity has been shown to increase with age in mouse and human tissues^14,15^. For example, canonical p53 target genes were upregulated in physiologically aged human fibroblasts^40–42^ (Extended Fig. 1a, b). To extend this, we investigated p53 activity at multiple ages, from young adult, through aged, to geriatric, by combined single cell and spatial approaches. To do this, we profiled liver from male and female mice at young (3-5 mo), old (21-24 mo) and geriatric (29-31 mo) ages using 10X multiome single-nucleus RNA-seq (snRNA-seq) and single-nucleus assay for transposase-accessible chromatin using sequencing (snATAC-seq). This approach identified the known major cell types in liver (Fig. 1a). Because hepatocytes are the principal parenchymal cell of the liver and carry out most liver-specific metabolic functions, we focused subsequent analyses on this population. A p53 target gene module score, based on the gene list of Fischer^43^ (Supplementary Table 1, p53 target genes), increased significantly with age in hepatocytes (Fig. 1b). Differentially expressed gene (DEG) analysis between geriatric and young hepatocytes revealed numerous canonical p53 target genes upregulated with age, including *Cdkn1a* and *Gadd45a*, both of which were upregulated at the RNA and protein level (Fig. 1c–f). To confirm these results *in situ*, we performed spatial transcriptomics using the Vizgen MERSCOPE platform^44^ with a custom 300-gene multiplexed error-robust fluorescence in situ hybridization (MERFISH) panel (Supplementary Table 1, SenNet MERFISH panel) comprising senescence, p53 target genes, inflammation, ISGs, and cell-type identification markers. Across all ages, p53-high hepatocytes defined by top 5% signaling, were preferentially found near other p53-high hepatocytes (Extended Fig. 1c), suggesting that high-p53 hepatocytes occupy spatially distinct microenvironments in aged liver. Confirming the 10x multiome analysis, MERSCOPE showed that p53 target gene score similarly increased with age in hepatocytes (Fig. 1g, Extended Fig. 1d). Together, these results show that expression of p53 target genes increases in aged mouse hepatocytes, most markedly in geriatric mice.

**Figure 1:**
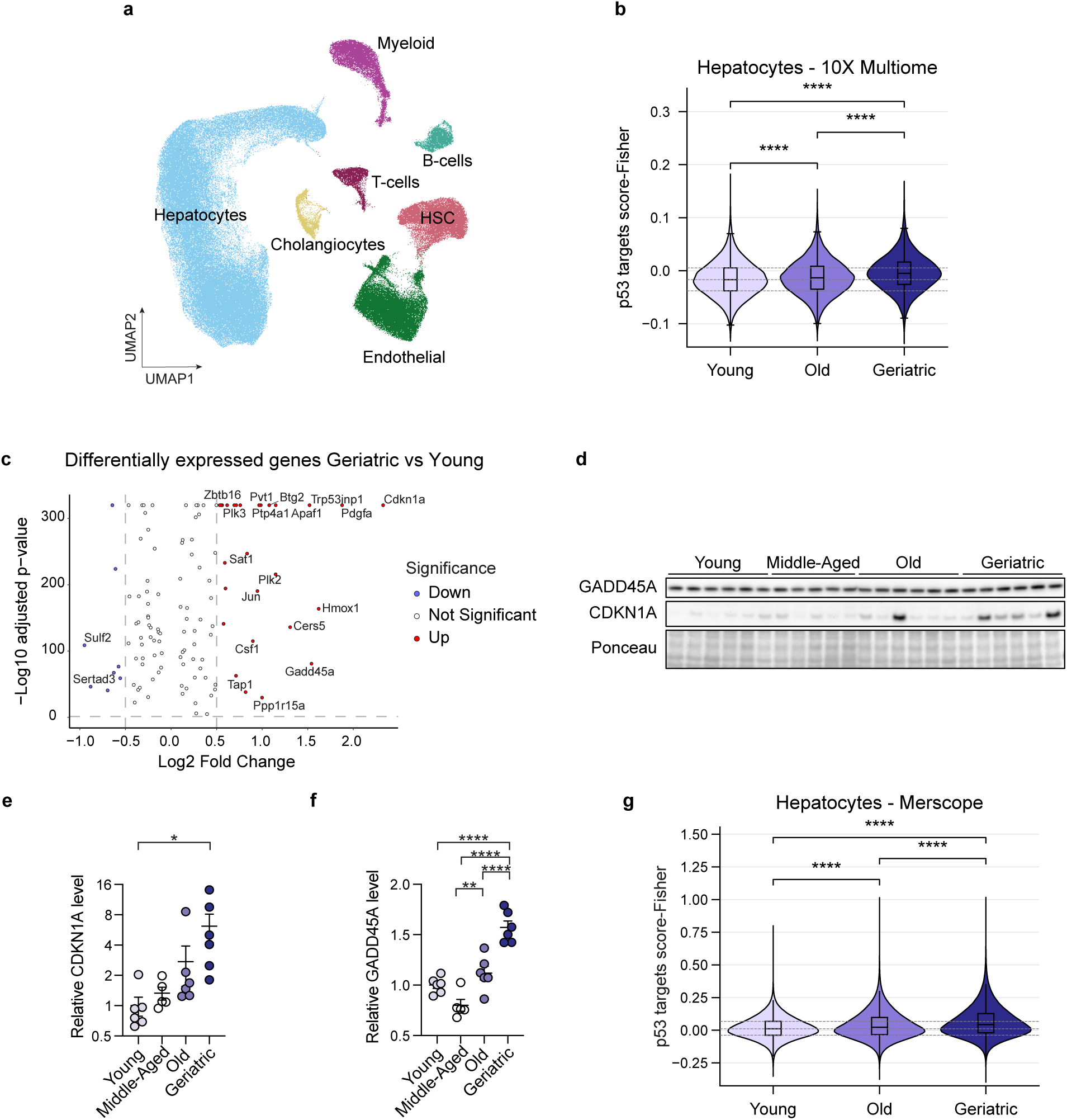
p53 activity increases in aged hepatocytes. a, UMAP representing 10X multiome profiling of liver, n = 6 mice per group. **b,** Per-cell p53 target module score in hepatocytes by age calculated from the 10X multiome dataset (Young, n = 38,110 cells from 6 mice; Old, n = 30,215 cells from 6 mice; Geriatric, n = 22,860 cells from 6 mice). *P* values: young vs old, *P* = 4 × 10^-33^; old vs geriatric, *P* = 9.7 × 10^-189^; young vs geriatric, *P* < 1 × 10^-300^. **c,** Differentially expressed p53 target genes in hepatocytes between geriatric and young mice, n = 6 mice per group. **d,** Western blot of CDKN1A and GADD45A, n = 6 mice per group and n = 5 for middle-aged. Ponceau was used as loading control. **e, f,** Quantification of western blot. **e,** CDKN1A, *P* values: young vs geriatric, *P* = 0.0265; middle-aged vs geriatric, *P* = 0.0529. **f,** GADD45A, *P* values: young vs geriatric, *P* < 0.0001; middle-aged vs geriatric, *P* < 0.0001; old vs geriatric, *P* < 0.0001; middle-aged vs old, *P* = 0.0059. **g,** Per-cell p53 target module score in hepatocytes by age calculated from the MERFISH dataset (Young, n = 417,887 cells from 4 mice; Old, n = 539,927 cells from 4 mice; Geriatric, n = 557,385 cells from 6 mice). *P* values: young vs old, *P* < 1 × 10^-300^; old vs geriatric, *P* < 1 × 10^-300^; young vs geriatric, *P* < 1 × 10^-300^. (b, g) Violins show the kernel density estimate of single-cell scores; inner boxplot line shows the median; box shows the 25th–75th percentile; whiskers extend to 1.5 × IQR. Dashed lines mark the 25th percentile, median and 75th percentile of the young distribution. (b, g) Statistical significance was assessed with two-sided Mann–Whitney *U* test on per-cell scores. (e, f) Statistical significance was assessed with one-way ANOVA with Tukey’s correction for multiple comparisons. Data shown as means ±SEM. \*\**P* < 0.01, \*\*\*\**P* < 0.0001.

### p53 suppresses aberrant cell-cycle activation and replication stress in aged mouse hepatocytes

Because of the age-dependent increase in p53 target gene expression, we set out to ask whether hepatocytes have an age-dependent response to loss of p53. To accomplish this, we used a single adeno-associated virus (AAV)-mediated CRISPR system expressing *Staphylococcus aureus* Cas9 (saCas9) under the hepatocyte-specific thyroxine-binding globulin (TBG) promoter, together with a U6-driven single-guide RNA (sgRNA)^45^. Mice received either an sgRNA targeting *Trp53* (sgTrp53) or an sgRNA targeting the safe-harbor locus *Rosa26* (sgRosa26) as a control. Western blot analysis of isolated hepatocytes confirmed efficient loss of p53 protein following sgTrp53 delivery (Fig. 2a). To determine whether the effects of p53 loss differ with age, we performed hepatocyte-specific KO in young (4 mo) and old (21 mo) mice and analyzed the livers 12 weeks later (Fig. 2b). The DNA editing efficiency was lower in old mice whole-livers, potentially due to the reported reduction of hepatocytes in aged liver tissue^46^ (Fig. 2c, Extended Fig. 2a,b). Bulk RNA-seq of whole liver confirmed downregulation of canonical p53 target genes in sgTrp53-infected old livers (Fig. 2d).

**Figure 2:**
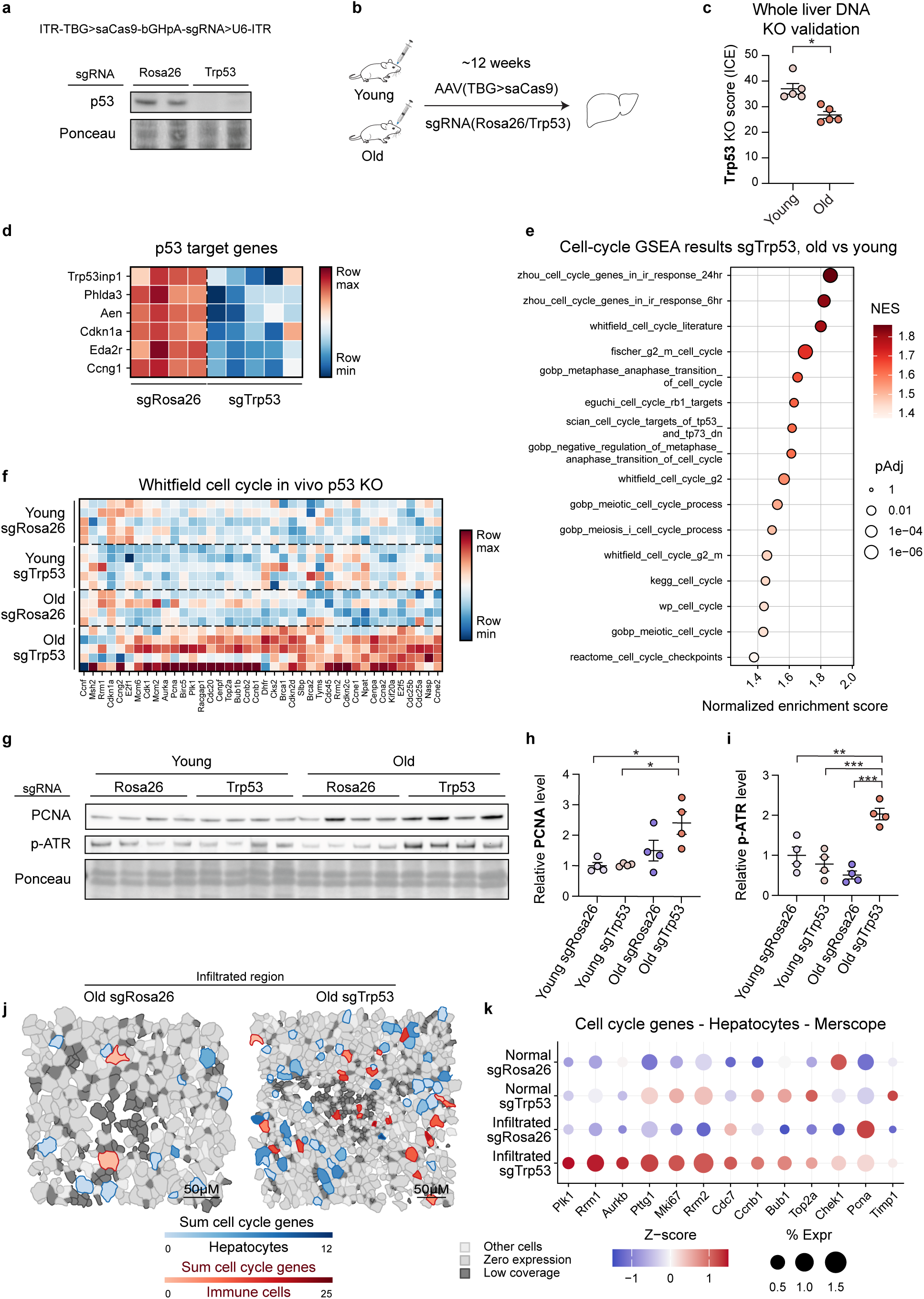
p53 suppresses aberrant cell-cycle activation and replication stress in aged mouse hepatocytes. a, Schematic representation of the AAV construct and western blot for p53 of isolated hepatocytes after treatment with either sgRosa26 or sgTrp53. **b,** Schematic representation of the experimental design to compare the effect of loss of p53 in young and aged liver hepatocytes (Young sgRosa26, n = 5; Young sgTrp53, n = 5; Old sgRosa26, n = 4; Old sgTrp53, n = 5). **c,** Quantification of the KO score in young and aged whole mouse liver using Inference of CRISPR Edits (ICE) analysis. *P* values: young vs old, *P* = 0.0079. **d,** Heatmap of canonical p53 target genes from bulk RNA-seq data. **e,** GSEA of cell-cycle pathways comparing young and aged sgTrp53-treated mice. **f,** Heatmap of Whitfield cell-cycle genes from bulk RNA-seq data. **g,** Western blot of PCNA and p-ATR, n = 4 per age per sgRNA treatment. Ponceau was used as loading control. **h, i,** Quantification of western blot. **h,** PCNA, *P* values: young sgRosa26 vs old sgTrp53, *P* = 0.0101; young sgTrp53 vs old sgTrp53, *P* = 0.0119. **i,** p-ATR, *P* values: young sgRosa26 vs old sgTrp53, *P* = 0.0035; young sgTrp53 vs old sgTrp53, *P* = 0.0007; old sgRosa26 vs old sgTrp53, *P* = 0.0001. **j,** Cell segmentation maps of cell-cycle gene expression in hepatocytes (blue gradient) and immune cells (red gradient). Other cell types are shown as light grey, zero-expressing cells as medium grey, and low-coverage cells as dark grey. Scale bar, 50 µm. **k,** Dot plot of cell-cycle genes in hepatocytes from infiltrated regions of sgRNA-treated livers; livers were profiled using a MERFISH spatial transcriptomics panel. n = 3 mice per sgRNA treatment; Normal sgRosa26, n = 15,091 cells; Normal sgTrp53, n = 7,798 cells; Infiltrated sgRosa26, n = 3,563 cells; Infiltrated sgTrp53, n = 4,519 cells. (c) Statistical significance was assessed with two-sided Mann–Whitney *U* test on per-mouse values. (h, i) Statistical significance was assessed with one-way ANOVA with Tukey’s correction for multiple comparisons. Data shown as means ±SEM. \**P* < 0.05, \*\**P* < 0.01, \*\*\**P* < 0.001.

A canonical function of p53 is to restrain the cell cycle, for example in response to DNA damage. Consistent with this, irradiated IMR90 cells depleted of p53 by siRNA transfection 4 days after irradiation (IR) showed decreased expression of *TP53* and its target gene *CDKN1A* (Extended Fig. 2c, d), and increased frequency of S-phase cells, as measured by 5-ethynyl-2′-deoxyuridine (EdU) incorporation (Extended Fig. 2e, f). In addition, bulk RNA-seq revealed broad upregulation of cell-cycle genes relative to IR siNTC controls (Extended Fig. 2g)^47^. At the protein level, the DNA replication stress marker phospho-ATR was also significantly elevated (Extended Fig. 2h). Remarkably, inactivation of p53 in old mouse liver induced a similar response. Gene set enrichment analysis (GSEA)^48^ of young versus aged sgTrp53 livers revealed selective enrichment of numerous cell-cycle gene sets in aged sgTrp53 liver (Fig. 2e). This was also evident in a heatmap of cell-cycle genes^47^, whose expression increased in aged sgTrp53 livers, compared to aged sgRosa26 and young sgTrp53 livers (Fig. 2f). Consistent with this transcriptional program, Proliferating Cell Nuclear Antigen (PCNA) protein was elevated in aged sgTrp53 livers (Fig. 2g, h). Significantly, phospho-ATR was also strongly increased specifically in aged sgTrp53 livers, compared to young sgTrp53 and aged sgRosa26 controls, suggesting replication stress in aged sgTrp53 hepatocytes (Fig. 2g, i).

To further validate the molecular effects of loss of p53 at the single-cell level, we profiled livers using complementary dissociative and spatial technologies. First, we performed 10X multiome snRNA-seq on young and old sgTrp53-treated livers, recovering the main hepatic cell types (Extended Fig. 3a). In old sgTrp53 hepatocytes, numerous cell-cycle genes were upregulated at the single-cell level (Extended Fig. 3b). Second, we used a custom MERFISH panel (Supplementary Table 1, p53 KO old MERSCOPE panel) targeting liver and immune cell identity, cancer and aging genes, including ISGs, to compare aged sgRosa26- and sgTrp53-treated liver. Since aging is typically accompanied by immune infiltration, we separately analyzed immune infiltrated and non-infiltrated tissue regions. Supporting the multiome results, hepatocytes in sgTrp53-treated regions showed an upregulation of cell-cycle genes, particularly in immune infiltrated regions (Fig. 2j, k). Taken together, these results show that inactivation of p53 in hepatocytes of old, but not young, mouse liver drives a transcriptional program indicative of at least partial cell-cycle entry, together with markers of DNA replication and replication stress.

### Loss of p53 elicits an age-specific transcriptional response and drives inflammation in aged liver

Having shown an age-dependent cell cycle response to loss of p53, we next performed an unbiased analysis of the transcriptomic response to p53 inactivation. Clustering of DEGs followed by Gene Ontology (GO) enrichment analysis identified five clusters of genes reflecting an age-dependent response to p53 loss (Fig. 3a; cluster gene lists in Supplementary Table 2). Of these clusters, Cluster 1 consisted of DEGs upregulated in both young and old sgTrp53-treated livers and showed terms related to circadian rhythm (Extended Fig. 4a); Cluster 2 consisted of DEGs upregulated only in young sgTrp53 livers, including genes involved in vitamin transmembrane transport, nuclear migration, and import across plasma membrane (Extended Fig. 4b); and Cluster 3 consisted of DEGs downregulated in both young and aged sgTrp53 livers, including genes related to heat response. However, most of the DEGs fell into Clusters 4 and 5. Cluster 4 consisted of DEGs downregulated only in aged sgTrp53 livers, reflecting small molecule catabolic processes and carboxylic acid catabolic process (Extended Fig. 4c). Cluster 5 consisted of DEGs upregulated only in aged sgTrp53 livers, representing an increase in many immune- and inflammation-associated genes (Extended Fig. 4d). Similarly, unbiased clustering of DEGs after p53 knockdown (KD) in irradiated IMR90 cells identified downregulation of metabolic genes (Cluster 2) and upregulation of pro-inflammatory pathways (Cluster 6) (Extended Fig. 4e; cluster gene lists and GO biological process (BP) terms in Supplementary Table 3). These results indicate that the consequences of p53 inactivation in hepatocytes of young and old liver are markedly different; inactivation of p53 specifically in aged hepatocytes causes marked changes in the immune/inflammatory and metabolic transcriptomes of whole liver.

**Figure 3:**
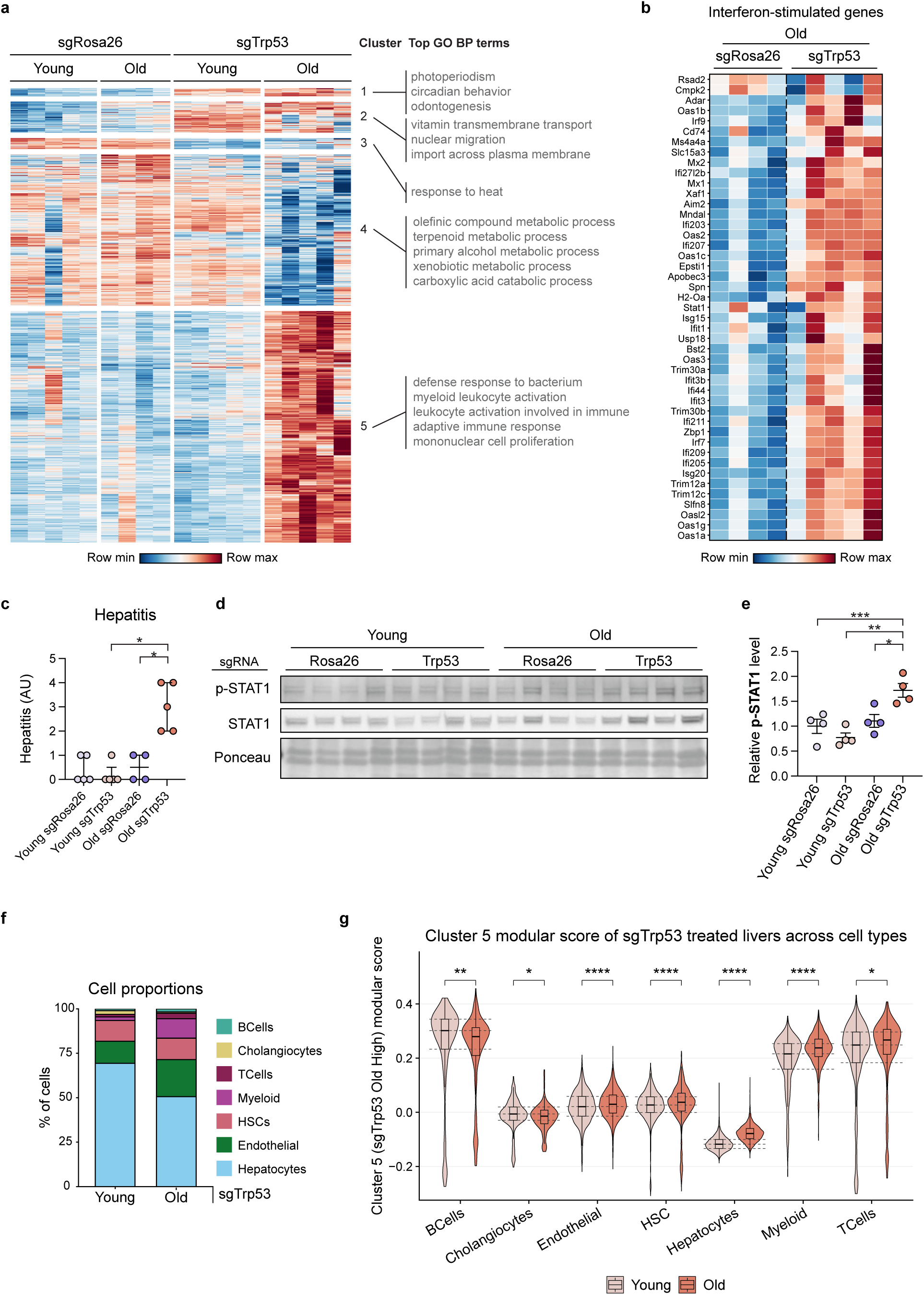
Loss of p53 elicits an age-specific transcriptional response and drives inflammation in aged liver. a, Heatmap of the DEGs in the following comparisons: young sgRosa26 vs young sgTrp53, young sgTrp53 vs old sgTrp53, old sgRosa26 vs old sgTrp53. In addition, the top 3 GO BP terms are shown. **b,** Heatmap of canonical ISGs in old mice treated with either sgRosa26 or sgTrp53 (Old sgRosa26, n = 4; Old sgTrp53, n = 5). **c,** Quantification of liver hepatitis by a board-certified pathologist blinded to age and treatment (Young sgRosa26, n = 5; Young sgTrp53, n = 5; Old sgRosa26, n = 4; Old sgTrp53, n = 5). *P* values: young sgTrp53 vs old sgTrp53, *P* = 0.032; old sgRosa26 vs old sgTrp53, *P* = 0.048; age × sgRNA interaction, *P* = 0.005. **d,** Western blot of STAT1 and p-STAT1, n = 4 per age per sgRNA treatment. Ponceau was used as loading control. **e,** Quantification of western blot, relative p-STAT1 level. *P* values: young sgRosa26 vs old sgTrp53, *P* = 0.008; young sgTrp53 vs old sgTrp53, *P* = 0.001; old sgRosa26 vs old sgTrp53, *P* = 0.0230. **f,** Bar plot representing the cell types present in sgTrp53-treated livers, n = 2 per group. **g,** Per-cell cluster 5 module score across cell types from the 10X multiome dataset of young and aged sgTrp53-treated livers (Young sgTrp53, n = 15,273 cells from 2 mice; Old sgTrp53, n = 15,146 cells from 2 mice). For each cell type, the cluster 5 gene set was restricted to genes detected in at least 5% of the young cells of that cell type before scoring with Seurat AddModuleScore. *P* values were calculated by young sgTrp53 vs old sgTrp53, two-sided Mann–Whitney *U* test, Benjamini–Hochberg-adjusted across cell types: B cells, *P* = 2.5 × 10^-3^; cholangiocytes, *P* = 2.3 × 10^-2^; endothelial, *P* = 8.5 × 10^-9^; HSC, *P* = 6.2 × 10^-8^; hepatocytes, *P* < 1 × 10^-300^; myeloid, *P* = 9.1 × 10^-13^; T cells, *P* = 2.1 × 10^-2^. (g) Violins show the kernel density estimate of single-cell scores; inner boxplot line shows the median; box shows the 25th–75th percentile; whiskers extend to 1.5 × IQR. Dashed lines mark the 25th, 50th and 75th percentiles of the young cells of each cell type. (c) Statistical significance was assessed with two-tailed exact Mann–Whitney *U* tests with Holm–Bonferroni correction across four pre-specified comparisons, and the age × sgRNA interaction with an exact permutation test; data shown as medians, each point is one mouse. (e) Statistical significance was assessed with one-way ANOVA with Tukey’s correction for multiple comparisons. Data shown as means ±SEM. \**P* < 0.05, \*\**P* < 0.01, \*\*\**P* < 0.001, \*\*\*\**P* < 0.0001.

First, we set out to validate and investigate the age-specific immune/inflammatory genes, whose expression is repressed by p53 specifically in aged liver (Cluster 5). Heatmaps of ISGs and inflammatory signaling genes showed their overall upregulation in whole liver of aged sgTrp53-targeted animals relative to aged sgRosa26 controls (Fig. 3b, Extended Fig. 5a). A heatmap of ISGs and hallmark inflammatory response genes also showed coherent upregulation in irradiated sip53-treated IMR90 cells relative to siNTC (Extended Fig. 5b, c). Consistent with the whole-liver transcriptome analysis, expert pathology assessment of liver H&E revealed increased foci of immune cell infiltration (hepatitis) in aged sgTrp53 livers relative to both aged sgRosa26 and young sgTrp53 controls (Fig. 3c, Extended Fig. 5d). Also consistent with the global increase in inflammatory gene expression, phospho-STAT1 was elevated in aged sgTrp53-treated livers by Western blot (Fig. 3d, e). Together, these results indicate that expression of p53 in hepatocytes restrains a pro-inflammatory program in aged liver.

Next, we set out to identify the cell types responsible for increased inflammatory gene expression (Cluster 5) in livers containing p53-deficient hepatocytes. To do this, we analyzed the 10X multiome snRNA-seq data from young and aged sgTrp53 livers. This analysis revealed age-associated shifts in cellular composition, with a reduction in the hepatocyte proportion and an increase in myeloid and T-cell fractions in aged sgTrp53 livers (Fig. 3f). Compared to young sgTrp53 livers, Cluster 5 genes were significantly and markedly upregulated in hepatocytes of aged sgTrp53 livers, the cell type targeted by TBG-driven KO (Fig. 3g). More modest but significant increases in Cluster 5 genes occurred in endothelial cells and hepatic stellate cells (HSCs) of aged sgTrp53 livers, and marked increases occurred in immune cells, specifically myeloid cells and T cells (Fig. 3g). These results show that the increased immune/inflammatory gene expression in old sgTrp53 that characterizes Cluster 5 is driven by increased gene expression in hepatocytes themselves, as well as myeloid cells and T cells. Our observations suggest that inactivation of p53 in aged hepatocytes drives inflammation and immune infiltration through communication with immune cells.

### Spatial transcriptomics reveals rewired hepatocyte–immune signaling in aged p53-deficient liver

To understand how aged sgTrp53 hepatocytes presumably communicate with infiltrating immune cells, we returned to our custom MERFISH panel and compared hepatocytes within immune-infiltrated regions to those in histologically normal regions of the same livers. Hepatocytes from infiltrated regions of aged sgTrp53 livers showed upregulation of several pro-inflammatory programs including antigen presentation (Cd74, H2-Aa, H2-Ab1, Lgmn, Tap1), interferon and innate immunity (Stat1, Gbp2, Zbp1, Ccl5, Plac8), phagocytosis and complement (Cd5l, Axl, C1qa, Sirpa), extracellular matrix (Col1a1, Col3a1), and stress response (Hmox1, Prkcd, Pkm), compared to non-infiltrated old sgTrp53 and infiltrated old sgRosa26 (Fig. 4a; Extended Fig. 6a). When quantified across all hepatocytes in infiltrated regions, summed major histocompatibility complex class II (MHC-II) and ISG expression levels were both significantly higher in old sgTrp53 than in old sgRosa26 hepatocytes (Fig. 4b). In addition, spatial mapping showed upregulation of MHC-II gene expression in aged sgTrp53 hepatocytes of infiltrated regions (Fig. 4c), consistent with previous reports that hepatocytes under stress can express MHC-II and promote the recruitment and activation of immune cells^49^. The same gene signatures were identified in Cluster 5 of bulk RNA-seq (Fig. 3a). Our results suggest that loss of p53 in aged hepatocytes activates an antigen-presentation and interferon program within the hepatocytes themselves, either of which may in turn facilitate immune infiltration.

**Figure 4:**
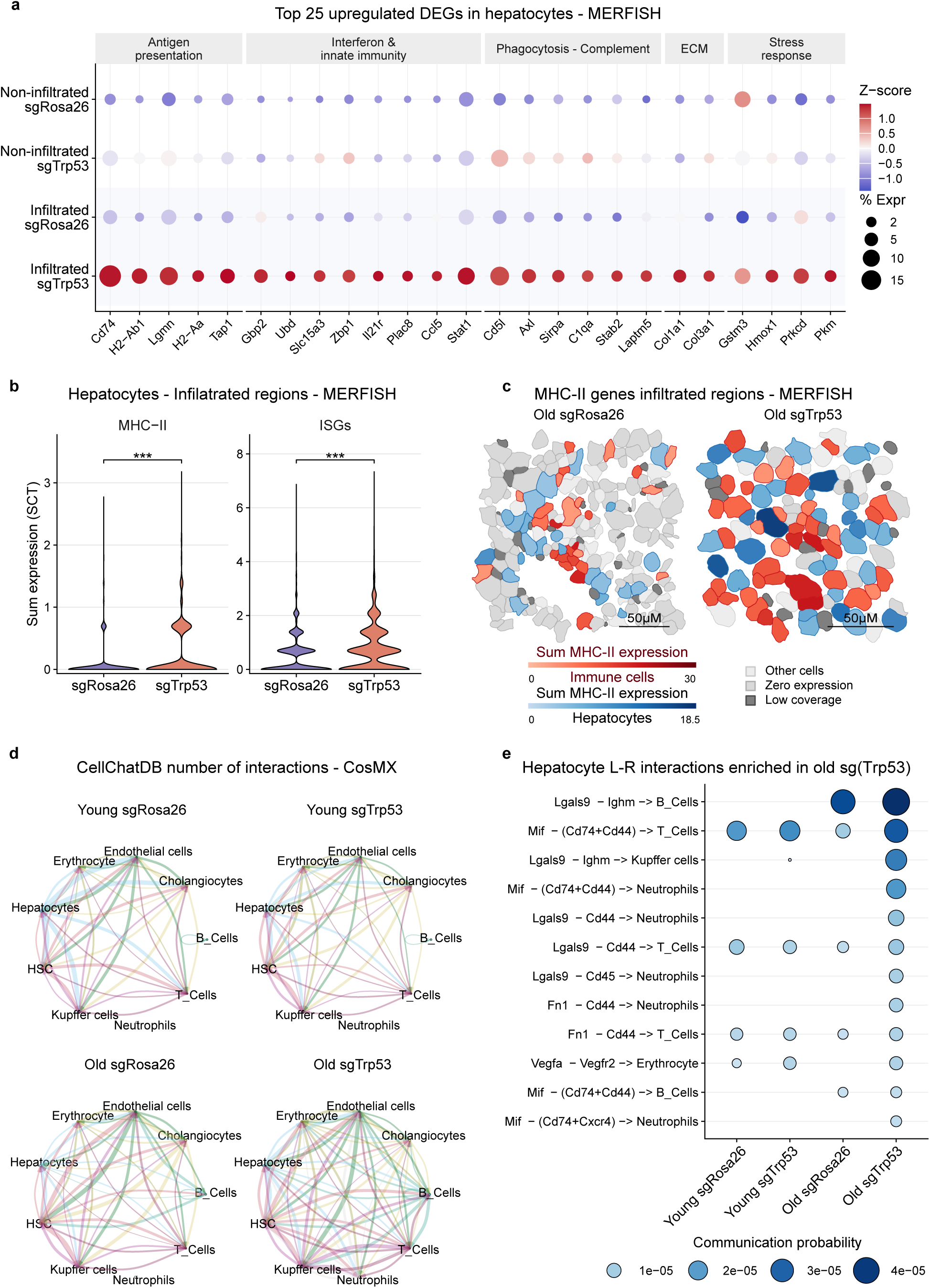
Spatial transcriptomics reveals rewired hepatocyte–immune signaling in aged p53-deficient liver. a, Dot plot of the top 25 concordant upregulated genes in hepatocytes from infiltrated regions of aged sgTrp53 treated livers; livers were profiled using a MERFISH spatial transcriptomics panel. n = 3 mice per sgRNA treatment; Normal sgRosa26, n = 15,091 cells; Normal sgTrp53, n = 7,798 cells; Infiltrated sgRosa26, n = 3,563 cells; Infiltrated sgTrp53, n = 4,519 cells. Genes were selected if significant in both comparisons infiltrated sgTrp53 versus infiltrated sgRosa26 hepatocytes and infiltrated versus normal hepatocytes within sgTrp53 livers; the 25 shown are ranked by *P* value and grouped by GO. Dot size represents the percentage of cells in each group expressing the gene; dot color represents the per-gene Z-score across the four group-mean expression values. **b,** Violin plots of summed MHC-II (Cd74, H2-Ab1, H2-Aa, H2-Eb1) and ISGs (28 genes; Supplementary Table 1, ISGs) expression per hepatocyte in infiltrated regions of aged sgRosa26 vs sgTrp53 treated livers. Values are calculated as sum of SCT log-normalized expression across the gene set per cell. *P* values were calculated by cell-level two-sided Wilcoxon rank-sum test. *P* values: MHC-II, *P* = 2.1 × 10^-107^; ISGs, *P* = 5.8 × 10^-42^. **c,** Cell segmentation maps of MHC-II gene expression in hepatocytes (blue gradient) and immune cells (red gradient). Other cell types are shown as light grey, zero expressing cells as medium grey, and low coverage cells as dark grey. Scale bar, 50 µm. **d,** Circular network plots of statistically significant cell-cell interactions (CellChatDB, *P* < 0.05). Pointed edges represent signaling from source to target cell type; line thickness is proportional to the number of significant L-R interactions between the two cell types. Line color matches the source cell type. **e,** Bubble plot of the top 12 hepatocyte-outgoing L-R interactions upregulated in old sgTrp53 livers (CellChatDB, *P* < 0.05; communication probability in old sgTrp53 greater than in each of the other three groups). Each row shows an L-R pair (source cell type → target cell type), ranked by communication probability in old sgTrp53. Dot size and dot color (blue gradient) both represent communication probability.

To further dissect the cell–cell interactions that transmit the consequences of sgTrp53 in aged hepatocytes to other cell types in the liver, we performed spatial transcriptomics using the CosMx Mouse Universal Cell Characterization RNA Panel^50^, which identified most liver cell types (Extended Fig. 6b). We observed a decrease in the proportion of hepatocytes and an increase in numerous immune cell types, including Kupffer cells, T cells, and B cells, in aged sgTrp53 mice compared to all other groups (Extended Fig. 6c–e). CellChatDB analysis^51^ of spatially adjacent cells revealed an increased number of interactions in aged sgTrp53 livers (Fig. 4d, Extended Fig. 6f). Focusing on interactions in which ligand expression increased in aged sgTrp53 hepatocytes by multiome (Extended Fig. 6g), we identified several candidate interactions enriched in aged sgTrp53 livers but absent or weak in young sgTrp53 and aged sgRosa26 controls, including hepatocyte *Lgals9* interaction with B cell and Kupffer cell *Ighm*, and hepatocyte *Mif* interaction with Cd74/Cd44 receptor on numerous immune cell types (Fig. 4e, Extended Fig. 6h). MIF is an inflammatory cytokine that promotes leukocyte recruitment^52^, while galectin-9 is an immunomodulatory lectin with context-dependent effects on immune cells^53^. In summary, these results indicate that in aged hepatocytes, p53 represses signals that are linked to liver inflammation and immune infiltration of the aged liver.

### p53 regulates metabolic genes in aged liver

In addition to suppressing immune infiltration and inflammation, bulk RNA-seq showed that p53 is required specifically in old liver for expression of many metabolic genes, especially those involved in lipid metabolism pathways, suggesting that p53 maintains metabolic homeostasis in aged liver (Cluster 4, Fig. 3a). We set out to define the cell types and mechanism through which p53 maintains expression of these genes in old liver. Analysis of snRNA-seq data showed that Cluster 4 metabolic genes are downregulated predominantly in aged sgTrp53 hepatocytes, consistent with TBG-directed hepatocyte-specific KO and the preferential expression of these genes in hepatocytes (Fig. 5a). GO analysis of all genes significantly downregulated in hepatocytes of old compared to young sgTrp53 mice confirmed a requirement for p53 in the expression of metabolic genes in aged hepatocytes, especially those involved in fatty acid and lipid metabolism (Extended Fig. 7a). To explore how p53 regulates the metabolic program of aged hepatocytes, we constructed a p53 gene regulatory network from our snATAC-seq data. We first identified open chromatin regions in young and aged hepatocytes and scanned all peaks genome-wide for p53 binding motifs using the JASPAR2020 database^54^. Each p53 motif-containing peak was assigned to its nearest annotated transcription start site (TSS; mm10), retaining peaks whose target gene was expressed in hepatocytes (mean normalized expression ≥ 0.1). We then integrated this set with DEGs from bulk RNA-seq of aged p53-KO livers, retaining only peak-linked genes whose expression changed upon p53 loss. This approach defined 194 candidate p53-activated (downregulated in old sgTrp53) and 45 candidate p53-repressed (upregulated in young sgTrp53) target genes in aged hepatocytes (Extended Fig. 7b). Strikingly, these genes were overwhelmingly enriched for metabolic ontologies, including catabolic processes (Fig. 5b). These integrated snRNA-seq and snATAC-seq results suggest that, in aged hepatocytes, p53 is directly involved in the expression of metabolic genes.

**Figure 5:**
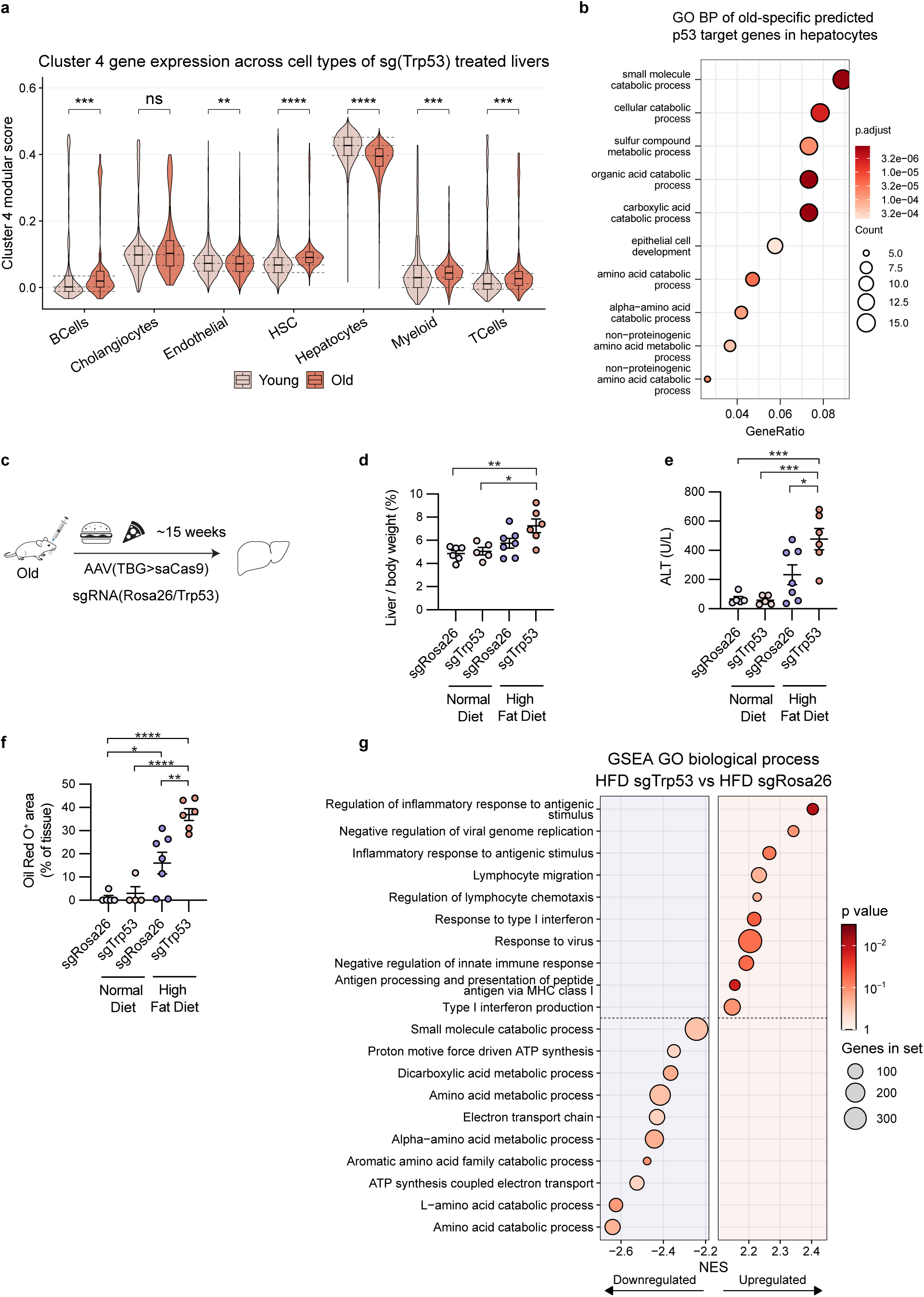
p53 regulates aged liver metabolism. a, Per-cell cluster 4 (from Fig. 3a) module score across cell types from the 10X multiome dataset from young and aged sgTrp53-treated livers (Young sgTrp53, n = 10,589 cells from 2 mice; Old sgTrp53, n = 7,664 cells from 2 mice). *P* values were calculated by young sgTrp53 vs old sgTrp53, two-sided Mann–Whitney *U* test, Benjamini–Hochberg-adjusted across cell types: B cells, *P* = 4 × 10^-4^; endothelial, *P* = 9 × 10^-3^; HSC, *P* = 1 × 10^-104^; hepatocytes, *P* < 1 × 10^-300^; myeloid, *P* = 1 × 10^-4^; T cells, *P* = 2 × 10^-4^. Cholangiocytes were not significant (ns). **b,** GO BP of old-specific predicted p53 target genes in hepatocytes. Over-representation against GO BP was tested with clusterProfiler; the top ten enriched terms are shown. Dot size indicates the number of genes per term; color indicates the adjusted *P* value; GeneRatio is calculated as the fraction of queried genes in each term. **c,** Schematic representation of the experiment designed to test the role of HFD in combination with sgTrp53 in aged mice. **d,** Liver-to-body-weight ratio after 15 weeks (normal diet sgRosa26, n = 6; normal diet sgTrp53, n = 5; HFD sgRosa26, n = 7; HFD sgTrp53, n = 6 mice). Group sizes differ across d–f owing to sample exclusion and availability; see Methods (High-fat diet) for details. *P* values: normal diet sgRosa26 vs HFD sgTrp53, *P* = 0.0044; normal diet sgTrp53 vs HFD sgTrp53, P = 0.0133. **e,** Serum ALT (n = 5, 5, 7 and 6 mice, in the order listed in d). P values: normal diet sgRosa26 vs HFD sgTrp53, P = 0.0005; normal diet sgTrp53 vs HFD sgTrp53, P = 0.0004; HFD sgRosa26 vs HFD sgTrp53, *P* = 0.0236. **f,** Positive area of Oil Red O staining per tissue area (n = 5, 4, 7 and 6 mice, in the order listed in d). *P* values: normal diet sgRosa26 vs HFD sgRosa26, P = 0.0281; normal diet sgRosa26 vs HFD sgTrp53, P < 0.0001; normal diet sgTrp53 vs HFD sgTrp53, P < 0.0001; HFD sgRosa26 vs HFD sgTrp53, P = 0.0013. **g,** GSEA of GO BP in HFD sgTrp53 vs HFD sgRosa26 livers (n = 6 and 7 mice). The ten most enriched terms in each direction are shown, ordered by NES; positive NES indicates enrichment in sgTrp53 HFD. Dot size indicates the number of genes per term and color indicates the nominal *P* value from rotation test. (a) Violins show the kernel density estimate of single-cell scores; inner boxplot line shows the median; box shows the 25th–75th percentile; whiskers extend to 1.5 × IQR. (d–g) Statistical significance was assessed with one-way ANOVA with Tukey’s correction for multiple comparisons. Data shown as means ±SEM. \**P* < 0.05, \*\**P* < 0.01, \*\*\**P* < 0.001, \*\*\*\**P* < 0.0001.

Accordingly, we wanted to test whether p53 is required to maintain metabolic homeostasis of the aged liver. To do this, we used the same AAV-CRISPR system to KO p53 (sgTrp53) or deliver a control sgRNA (sgRosa26) to livers of aged mice, then challenged both groups with high-fat diet (HFD) (Fig. 5c). Phenotypically, sgTrp53-treated mice on HFD showed a trend toward an elevated liver-to-body-weight ratio and significantly elevated serum alanine aminotransferase (ALT), a marker of liver damage, compared to HFD sgRosa26 controls (Fig. 5d, e). To test whether loss of p53 leads to impaired lipid handling, we quantified lipid accumulation by Oil-Red-O staining. Aged sgTrp53 livers on HFD showed larger Oil-Red-O^+^ area compared with HFD sgRosa26 livers (Fig. 5f; Extended Fig. 7c). RNA-seq of aged sgTrp53 and sgRosa26 livers on HFD identified similar p53-dependent programs as on normal diet, with GSEA showing upregulation of immune, inflammatory, type I interferon and antigen-presentation programs in sgTrp53-treated livers, together with downregulation of numerous catabolic pathways (Fig. 5g). Consistent with the elevated type I interferon signature, ISGs were broadly upregulated in aged sgTrp53 livers on HFD (Extended Fig. 7d). In line with its role in regulating metabolic gene programs, loss of p53 in aged hepatocytes increased susceptibility to dietary challenge, exacerbating hepatic lipid accumulation and inflammatory responses to HFD, indicating that hepatocyte p53 protects the aged liver against metabolic stress.

### Pharmacologic activation of p53 promotes features of healthy aging

These results suggest that pharmacologic activation of p53 in aged liver might dampen chronic inflammation and sustain metabolic function. Indeed, we previously showed that activation of p53 with the small-molecule MDM2 inhibitor HDM201 increases p53 levels in aged mouse liver, reverses age-associated transcriptomic changes in inflammation and lipid metabolism genes and attenuates age-associated immune infiltration, all measured in whole liver^34^. To define the effects of HDM201 at the single-cell level, we performed 10X multiome snRNA-seq and snATAC-seq on livers from aged mice treated with HDM201 or vehicle. Analysis identified all major liver cell types (Fig. 6a; Extended Fig. 8a). Immune cell proportions (myeloid, T and B cells) trended toward a decrease with HDM201 treatment (Extended Fig. 8b, c). We then analyzed module scores of the two age- and p53-dependent gene modules previously defined by bulk liver RNA-seq (Clusters 4 and 5, Fig. 3a) in snRNA-seq data. After HDM201 treatment, Cluster 5 (enriched for inflammatory genes) was significantly reduced in B cells, hepatocytes, myeloid cells and T cells, while Cluster 4 (enriched for fatty acid and lipid metabolism genes) was significantly upregulated in hepatocytes, myeloid cells and T cells (Fig. 6b). Consistent with this, pathway analysis of upregulated genes in hepatocytes from HDM201-treated mice showed numerous pathways related to fatty acid and lipid metabolism (Fig. 6c). Thus, pharmacologic activation of p53 in aged liver induces many cell type-specific gene expression changes reciprocal to hepatocyte-specific p53 KO, notably increased expression of metabolic genes in hepatocytes and decreased expression of inflammatory genes in hepatocytes, B cells, myeloid cells and T cells, supporting a role for p53 in maintaining cell and tissue homeostasis in the aged liver.

**Figure 6:**
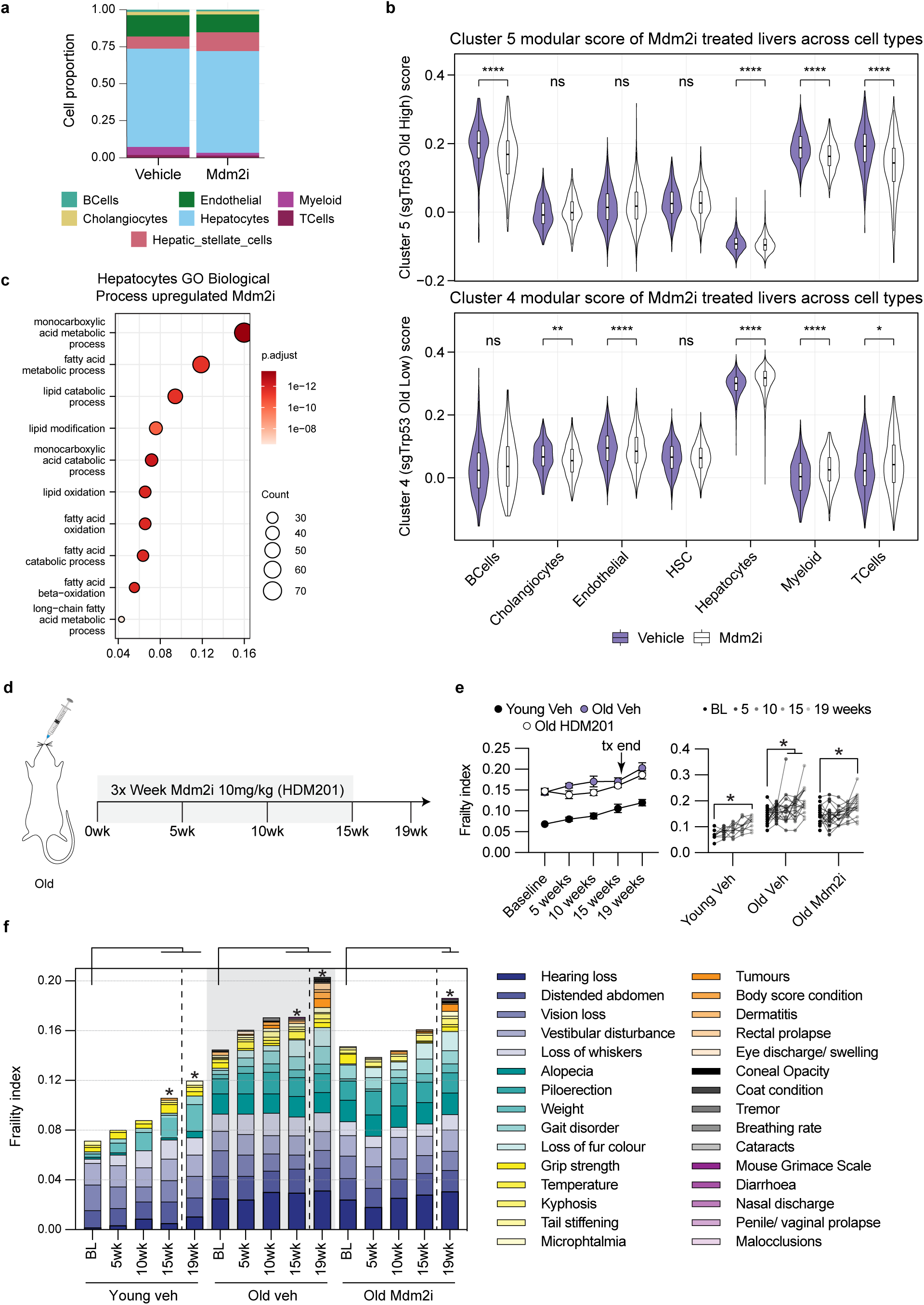
p53 promotes healthy aging. a, Bar plot representing the cell types of mice treated with HDM201 or vehicle control, n = 2 per group. **b,** Per-cell module scores of cluster 5 (top) and cluster 4 (bottom) across cell types from the snRNA-seq dataset of aged livers treated with vehicle or Mdm2i (Vehicle, n = 17,247 cells from 2 mice; Mdm2i, n = 17,205 cells from 2 mice). For each cell type, the gene set was restricted to genes detected in at least 5% of the vehicle cells of that cell type before scoring with Seurat AddModuleScore. *P* values were calculated by vehicle vs Mdm2i, two-sided Mann–Whitney *U* test, Benjamini–Hochberg-adjusted across cell types within each panel. Cluster 5: B cells, *P* = 1.8 × 10^-7^; hepatocytes, *P* = 7.6 × 10^-18^; myeloid, *P* = 7.6 × 10^-18^; T cells, *P* = 2.4 × 10^-14^; cholangiocytes, endothelial and HSC were not significant (ns). Cluster 4: cholangiocytes, *P* = 1.2 × 10^-3^; endothelial, *P* = 9.5 × 10^-5^; hepatocytes, *P* = 1.1 × 10^-309^; myeloid, *P* = 6.1 × 10^-8^; T cells, *P* = 3.5 × 10^-2^; B cells and HSC were not significant (ns). **c,** GO BP enrichment analysis of genes upregulated in hepatocytes of Mdm2i-vs vehicle-treated aged livers (snRNA-seq, n = 2 mice per group). The ten most significant terms are shown. The dot size and color represent gene number and statistical significance (Benjamini–Hochberg-adjusted *P),* respectively. **d,** Schematic representation of the HDM201 treatment regime in aged mice. Mice were treated 3 times a week for 15 weeks with 10 mg/kg of HDM201 or vehicle. **e,** Frailty index of vehicle-or Mdm2i-treated mice, shown as group means (left) and individual mice (right); lines connect the same mouse. *P* values (vs baseline of the same group): Young Veh 15 weeks, P = 0.0015; Young Veh 19 weeks, P < 0.0001; Old Veh 15 weeks, P = 0.024; Old Veh 19 weeks, P = 0.0008; Old Mdm2i 19 weeks, P = 0.0044; all other timepoints vs baseline were not significant. **f,** Breakdown of the frailty index by category; bar height equals the group mean frailty index. *P* values as in e. n = 20 mice per old group and n = 10 young vehicle mice at the outset of treatment; mice that died or were euthanized during the study (3 Old Veh, 2 Old Mdm2i) contributed all measurements obtained before death. (b) Violins show the kernel density estimate of single-cell scores; inner boxplot line shows the median; box shows the 25th–75th percentile; whiskers extend to 1.5 × IQR. (e, f) Statistical significance was assessed with a linear mixed model for repeated measures (group × timepoint, unstructured within-mouse covariance with timepoint- and group-specific variances) with Šídák’s correction for the four post-baseline timepoints per group. Data shown as means ± SEM. *P < 0.05, **P < 0.01, \*\*\**P* < 0.001, \*\*\*\**P* < 0.0001.

To test whether pharmacologic activation of p53 has systemic benefits beyond the liver, we performed an extended duration study measuring frailty as a readout. Specifically, we treated aged mice with HDM201 three times weekly for 15 weeks and assessed frailty over time using a validated mouse clinical frailty index^55^ (Fig. 6d). We found no evidence of HDM201 toxicity at 16 weeks of treatment, with complete blood counts (CBCs) comparable to age-matched vehicle controls (Extended Fig. 8d–i). Vehicle-treated young and aged mice both showed progressive increases in frailty over the treatment period. However, HDM201-treated aged mice showed a significant delay in frailty accumulation compared to vehicle-treated aged mice (Fig. 6e, f). Four weeks after treatment cessation (week 19), frailty in HDM201-treated mice approached vehicle-treated levels, indicating that sustained p53 activation is required to maintain this protection. Together, these data identify p53 as a druggable target for healthy aging, with pharmacological activation sufficient to delay onset of frailty in aged mice.

## Discussion

Our work suggests that p53 is required for healthy aging in mice. Deleting p53 in hepatocytes produced an age-dependent response in which the aged liver showed 1) accumulation of markers of cell-cycle activation and replication stress, 2) a cell-intrinsic and extrinsic inflammatory response with immune infiltration, and 3) reduced expression of hepatocyte metabolic genes. Pharmacologic activation of p53 with the MDM2 inhibitor HDM201 had the opposite effect and also delayed the onset of organismal frailty. At the tissue level, loss of p53 in aged hepatocytes was associated with hepatitis and loss of metabolic homeostasis, effects that are typically associated with tissue dysfunction and predisposition to disease^56^. p53 activity in aged tissues appears therefore to be an adaptive response that supports healthy aging.

p53 target gene expression increases with age across numerous tissues and mammalian species^14,15^, and likely reflects the accumulation of genotoxic and metabolic stresses over the lifespan^11^. Under the AP model, this age-associated activity may be detrimental, particularly late in life^11^. Consistent with this model, p53 promotes senescence entry^7,8,^ and mice with elevated p53 activity exhibit signs of accelerated aging and have a shortened lifespan^13,27^. However, other mouse models with elevated but physiologically controlled p53 activity exhibit enhanced tumor resistance without accelerated aging ^35–37^, challenging this interpretation. Thus, whether endogenous p53 activity is beneficial or detrimental in aged tissues remains unresolved. Critically, this question has not been directly addressed through loss-of-function experiments in aged animals, because whole-body p53 KO mice develop cancer and die early in life^38,39^.

We therefore performed p53 loss-of-function experiments in young and aged wild-type mice, preserving p53 function during development. Using this approach, we found an age-dependent increase in expression of cell-cycle genes in p53 KO. This was much more pronounced in old hepatocytes and accompanied by signs of DNA replication stress, suggesting that p53 signaling restrains aberrant cell-cycle activation in aged hepatocytes. Our results are consistent with a “precarious balance” between oncogenic and tumor-suppressive programs, whereby tumor suppressive pathways, such as p53, restrain concurrent activation of oncogenic pathways. We have previously reported this dual activation of oncogenic and tumor suppressor pathways in the aging liver^57^. Such opposing cellular programs have also been described during the benign-to-malignant transition in the pancreas^58^. Our findings extend this framework by providing functional evidence that p53 helps maintain this balance in aged hepatocytes.

The age-dependent response to p53 loss extended beyond cell-cycle regulation, with a substantially greater transcriptional response in aged than in young liver. This response was characterized in part by increased inflammation in the aged liver. To determine the cellular basis of this response, we performed snRNA-seq and spatial transcriptomic analyses. Our analyses suggested that increased inflammation was associated with both greater immune cell infiltration and activation of hepatocyte-intrinsic pro-inflammatory pathways in aged hepatocytes following p53 loss. Spatial transcriptomic analysis using CellChat identified increased cell–cell communication in aged sgTrp53 livers, suggesting several potential mechanisms of immune cell recruitment, including interactions between hepatocyte-derived Lgals9 and Ighm on B cells and Kupffer cells, and between hepatocyte-derived Mif and Cd74/Cd44 receptors on numerous immune cell types. In addition, analysis of immune-infiltrated regions in aged sgRosa26 and sgTrp53 livers identified ISGs, MHC class II genes, and antigen-presentation pathways^49^ among the most prominently enriched signatures following p53 loss. Although p53 has previously been reported to positively regulate interferon signaling^59,60^, our findings suggest an indirect mechanism whereby p53 loss in aged hepatocytes exacerbates replication stress, activating cGAS–STING and downstream type I interferon signaling^61–65^. Given that increased interferon signaling is a hallmark of aging across multiple tissues^15,57^, p53 loss may amplify this pre-existing inflammatory program. Our findings extend prior evidence suggesting that p53 suppresses the inflammatory response in mice, and that pharmacologic stabilization of p53 could attenuate acute inflammatory injury ^66,67^. We therefore propose that in hepatocytes p53 restrains liver inflammation and immune cell infiltration in aged liver.

Another age-dependent arm of p53 was maintenance of metabolic genes. There are nearly 10^6^ p53 half-sites in the mammalian genome, with ∼10^3^ high-confidence sites associated with putative target genes^68,69^, of which only ∼10^2^ are reproducible across cancer cell lines^43^; p53 target genes in non-cancer contexts remain poorly defined. By integrating snATAC-seq with p53 motif analysis^54^ and bulk RNA-seq of sgTrp53-treated livers, we identified 194 candidate p53-activated and 45 candidate p53-repressed target genes in aged hepatocytes, strongly enriched for catabolic and lipid-metabolic processes. These findings suggest that p53 maintains metabolic gene expression in the aged liver, consistent with earlier reports that p53 promotes lipid catabolism and limits anabolism^70–73^. Previous studies have shown that pharmacologic p53 activation with low-dose doxorubicin reduces steatosis and liver damage in diet-induced metabolic dysfunction-associated steatotic liver disease (MASLD) and steatohepatitis (MASH) models^74^. The loss-of-function literature is conflicting, with reduced p53 activity reported both to promote^75^ and to reduce^76,77^ diet-induced steatosis. Our experiments address p53 function directly in aged liver, where p53 loss in hepatocytes under HFD challenge increased lipid accumulation and raised serum ALT, a biomarker of liver damage. These effects suggest direct transcriptional control of metabolic genes. Increased p53 activity in aged tissues might therefore act as a guardian of cell identity^78,79^ and, in hepatocytes, of their metabolic function.

To test whether pharmacologic activation of p53 could ameliorate age-associated decline, we stabilized p53 in aged mice using the MDM2 inhibitor HDM201. At the single-nucleus level, we found reduced immune recruitment and lower pro-inflammatory signaling across multiple cell types, suggesting a broad anti-inflammatory effect of p53 activation. HDM201 also enhanced hepatocyte expression of metabolic genes. To assess effects at the whole-animal level, we used a validated mouse clinical frailty index^55^ and found that HDM201-treated mice developed signs of frailty later than controls. The intermittent, modest p53 activation achieved here differs from the pro-apoptotic activation sought in oncology^80^, suggesting that aging interventions may require a distinct dosing strategy and therapeutic window. MDM2 inhibitors have, however, already been tested clinically in age-associated disease. UBX0101, a p53/MDM2 interaction inhibitor developed as a senolytic, did not reduce patient-reported knee pain relative to placebo after a single intra-articular injection in a Phase 2 osteoarthritis trial (NCT04129944)^81^. This trial carried out by Unity Biotechnology evaluated local treatment for symptomatic osteoarthritis, and does not directly address whether intermittent systemic p53 activation can improve age-associated function at a wider level. In addition, little has been published on drug exposure, p53 signaling activation or the patient selection guidelines from this study. Translating our findings into a clinical setting would require defining dosing regimens that achieve the intended activation and duration of p53 activation. In addition, it would be necessary to establish pharmacodynamic markers of target engagement, identify suitable patient populations, and evaluate safety together with functional outcomes relevant to aging.

Our study has several limitations. First, our loss-of-function experiments were performed in a single tissue, so we cannot say whether p53 supports healthy aging in other organs. However, HDM201 delayed the onset of frailty, a whole-organism phenotype, suggesting that the benefit of p53 activation is not exclusive to the liver. Second, the AAV-TBG-saCas9 system KOs p53 in a hepatocyte-specific but mosaic manner, and the effects measured across the whole liver are therefore diluted by cells that retain p53. Third, the loss-of-function experiments used male mice, and the activation experiments used female mice, so sex-specific effects remain untested. Finally, the link between p53 loss and the transcriptional changes we observe remains correlative.

In conclusion, we define an age-specific role for p53 during liver aging, in which endogenous p53 restrains aberrant hepatocyte cell-cycle activation, metabolic dysfunction and inflammation. Pharmacologic activation of p53 delayed the onset of frailty in aged mice, arguing that, contrary to AP predictions, p53 activity in aged tissue is protective rather than degenerative, and identifying p53 as a candidate target for interventions to promote healthy aging in humans.

## Methods

### Mice and housing

All animal experiments were approved by the Institutional Animal Care and Use Committee of Sanford Burnham Prebys Medical Discovery Institute (AUF 22-010, 23-026, 25-011 and 26-008) and performed in accordance with NIH guidelines. C57BL/6JN mice were obtained from the National Institute on Aging Aged Rodent Colony and housed under specific pathogen-free conditions at 21–24 °C and 30–70% humidity on a 12 h light/dark cycle with ad libitum access to food (Teklad 2018) and water. Young (3–5 months), middle-aged (12 months), old (21–24 months) and geriatric (29–31 months) mice were used as indicated in the figure legends. Mice were allowed to acclimate to the facility for 2 weeks on arrival before any procedure. Loss-of-function (AAV-CRISPR) experiments used male mice; MDM2 inhibitor experiments used female mice. Mice were randomly assigned to treatment groups within each age, and investigators performing histological and pathological scoring were blinded to age and treatment. Animals were monitored daily, weighed at least weekly during treatment, and euthanized by CO_2_ asphyxiation in the morning at the timepoints indicated, after which tissues were flash frozen.

## AAV-mediated hepatocyte-specific Trp53 KO

Hepatocyte-specific gene editing was achieved with a single AAV vector encoding Staphylococcus aureus Cas9 (saCas9) under the hepatocyte-specific TBG promoter together with a U6-driven sgRNA^45^. Guide RNAs were designed with CRISPick against the mouse reference genome GRCm38 (Ensembl v.102). The sgRNA targeting Trp53 (sgTrp53, 5’-GACAAAAGATGACAGGGGCCA-3’) and the control sgRNA targeting the Rosa26 safe-harbor locus (sgRosa26, 5’-CTCGATGGAAAATACTCCGAG-3’) were cloned into pX602-AAV-TBG::NLS-SaCas9-NLS-HA-OLLAS-bGHpA;U6::BsaI-sgRNA (Addgene #61593) and transformed into NEB Stable competent E. coli. Insertion was validated by direct colony sequencing (Genewiz) using a U6 promoter primer (5’-GACTATCATATGCTTACCGT-3’).

Hepatocyte-specific KO was performed in young (4 month) and old (21 month) mice (young sgRosa26, n = 5; young sgTrp53, n = 5; old sgRosa26, n = 4; old sgTrp53, n = 5). Mice received 3.33 × 10^11^ genome copies per mouse in 100 µL sterile saline by retro-orbital injection. Livers were harvested 12 weeks after injection, or 15 weeks after injection for the high-fat diet experiment.

## AAV production

Large-scale rAAV production was performed by the Sanford Burnham Prebys Functional Genomics Core Facility. In short, HEK-293T cells were co-transfected with the rAAV vector, AAV8 RepCap and Helper (Cell Biolabs) plasmid DNA using polyethylenimine (Polysciences). Ninety-six hours after transfection, crude virus was isolated from the supernatant by polyethylene glycol precipitation and from the cell pellet by sonication. Concentrated viral particles were purified by iodixanol (Millipore Sigma) gradient ultracentrifugation. The fraction containing rAAV particles was collected and buffer-exchanged into PBS before being aliquoted and stored at -80 °C. Viral titer was determined by SYBR Green qPCR.

## High-fat diet

Aged (21 month) male mice were injected with AAV-sgTrp53 or AAV-sgRosa26 as above and, on the same day, placed on a high-fat diet (HFD; TestDiet 58Y1, 5.10 kcal/g, 61.6% kcal from fat, 20.3% kcal from carbohydrate, 18.1% kcal from protein) or maintained on standard chow (Teklad 2018) for 15 weeks. Four groups were compared: normal diet sgRosa26 (n = 6), normal diet sgTrp53 (n = 6), HFD sgRosa26 (n = 7) and HFD sgTrp53 (n = 6). One normal diet sgTrp53 mouse showed no detectable Trp53 editing by Inference of CRISPR Edits (ICE) analysis and was excluded from all analyses, giving final group sizes of n = 6, 5, 7 and 6, respectively. Body weight was recorded weekly. At endpoint, blood was collected by cardiac puncture for serum chemistry, and ALT was measured as described under Blood profiling; serum was unavailable for one normal diet sgRosa26 mouse (n = 5, 5, 7 and 6). Final body weight and liver weight were recorded at takedown and used to calculate the liver-to-body-weight ratio. Livers were divided for flash freezing, OCT embedding and fixation in neutral-buffered formalin. OCT-embedded tissue was not available for one mouse in each normal diet group; Oil Red O quantification was therefore performed on n = 5, 4, 7 and 6.

## HDM201 (MDM2 inhibitor) treatment

HDM201 (siremadlin; Novartis) was suspended according to the manufacturer’s instructions in phosphate-buffered methylcellulose (0.5% methylcellulose in 50 mM phosphate buffer, pH 6.8) and administered by oral gavage at 10 mg/kg in the morning; control mice received vehicle on the same schedule. Two independent cohorts were used. The single-nucleus multiome analysis was performed on livers from the same aged female mice described previously^34^, which received HDM201 or vehicle daily for 14 days and were collected in the morning after the final dose. For the longitudinal frailty cohort, 18-month-old female C57BL/6JN mice were treated three times weekly (Monday, Wednesday and Friday) at the same dose for 15 weeks, and frailty was assessed at baseline and at weeks 5, 10, 15 and 19 (approximately 4 weeks after treatment cessation). The cohort comprised n = 20 mice per aged group and n = 10 young vehicle mice (4–5-month-old females) at the outset of treatment. No mice were excluded; mice that died or were euthanized at humane endpoints during the study (3 old vehicle, 2 old HDM201) contributed all measurements obtained before death.

## Frailty index

Frailty was assessed using a validated mouse clinical frailty index^55^ comprising numerous non-invasive items spanning the integument, musculoskeletal, vestibulocochlear/auditory, ocular/nasal, digestive/urogenital and respiratory systems, together with body surface temperature. Each item was scored 0 (absent), 0.5 (mild) or 1 (severe) and the frailty index was calculated as the sum of scores divided by the number of items assessed. Mice were scored at baseline and at weeks 5, 10, 15 and 19 by the same blinded investigator (R.A.).

## Blood profiling

Blood was collected at the timepoints indicated in the figure legends. For complete blood counts (CBCs), whole blood was collected retro-orbitally into BD Microtainer K2EDTA tubes (BD, cat# 365974) and analyzed on a VetScan HM5 hematology analyzer (Zoetis), reporting hemoglobin (HGB), red blood cells (RBC), platelets (PLT), white blood cells (WBC), lymphocytes (LYM), monocytes (MON) and neutrophils (NEU).

For serum chemistry, blood was collected retro-orbitally or by cardiac puncture, depending on the experiment, into Microvette CAT serum-gel tubes (Sarstedt), allowed to clot for 30 min at room temperature and centrifuged at 10,000 g for 10 min. The serum supernatant was transferred to a fresh tube, diluted 1:1 in saline and stored at −20 °C until analysis. Alanine aminotransferase (ALT) and other liver analytes were measured on a VetScan VS2 chemistry analyzer using VetScan Mammalian Liver Profile reagent rotors (Zoetis) according to the manufacturer’s instructions.

## Hepatocyte isolation

Primary hepatocytes were isolated by two-step collagenase perfusion, adapted from Charni-Natan et al.^82^ Mice were euthanized by CO_2_ asphyxiation and a 24 G × 3/4-inch catheter (Surflo, SR-0X2419CA) was inserted into the inferior vena cava below the liver. The liver was perfused with 50 mL of Hank’s Balanced Salt Solution (HBSS; Gibco, ref. 14175-095) pre-warmed to 42 °C at 6 mL min−1, followed by 40 mL of DMEM (Gibco, ref. 10313-021) containing collagenase type IV (Gibco, ref. 17104-019), also pre-warmed to 42 °C. After digestion, livers were excised and dissociated in ice-cold DMEM containing 2% bovine serum albumin (BioWorld, CAS 9048-46-8), filtered through a 100 µm cell strainer and kept on ice. Cell suspensions were centrifuged at 50 g for 2 min at 4 °C and washed five times in cold DMEM to remove debris. The pellet was resuspended in 40% Percoll (Cytiva, ref. 17089102) made up with 10% 10× HBSS (Gibco, ref. 14065-056) and 60% DMEM, and centrifuged at 100 g for 7 min at 4 °C to enrich viable hepatocytes. The resulting pellet was washed once more in DMEM at 50 g and resuspended. Viability and cell counts were determined by trypan blue staining using a hemocytometer. Hepatocytes were plated on collagen-coated 6-well plates at 2.5 × 10^5^ cells per well in DMEM supplemented with 10% FBS, 1% penicillin-streptomycin and 1% L-glutamine. The following morning, MG132 (dissolved in DMSO to a 50 mM stock) was added to a final concentration of 10 µM (1:5,000 dilution in culture medium), and cells were collected 24 h later for p53 immunoblotting.

## Genomic DNA extraction, Sanger sequencing and ICE analysis

KO was confirmed in mouse genomic DNA extracted from whole liver using the Monarch Genomic DNA Purification kit (New England Biolabs, ref T3010S), followed by PCR amplification of the ∼500 bp region surrounding the sgRNA target site. PCR was performed with 2X Taq PCR Master Mix (APExBIO, ref K1034), 100 ng of genomic DNA and 1 µl of 100 µM primers (Trp53 locus: forward 5’-GCTCTTGTTTTCCAGACTTCCTC-3’, reverse 5 GAGAAGAGGCTAAAAAGGTTCAGG-3’ Rosa26 locus: forward 5’-CTAGAAGATGGGCGGGAGTCT-3’, reverse 5’-TCTAGGGGTTGGATAAGCCAGT-3’) using an annealing temperature of 60 °C for 35 cycles. PCR products were purified with the QIAquick PCR Purification Kit (QIAGEN, ref 28104) and Sanger sequenced with the forward amplification primer (Genewiz). To quantify cut efficiency, ICE analysis was performed as described (ICE v3.0, Synthego; accessed 04/15/2024)^83^, with the corresponding uninjected or sgRosa26 sample as reference.

## Western blotting

Mouse liver tissue was lysed in modified RIPA buffer (Thermo Fisher Scientific, cat# 89900) using a Precellys tissue disruptor (Bertin Technologies); isolated hepatocytes and IMR90 cells were lysed in the same buffer without disruption. Lysis buffer was supplemented with 1x Halt protease and phosphatase inhibitor cocktail (Thermo Fisher Scientific, cat# 78440), and protein concentration was determined by Bradford assay (Thermo Fisher Scientific, cat# 23246). Equal amounts of protein (50 µg unless otherwise stated in the figure legend) were resolved on NuPAGE 4–12% Bis-Tris gels and transferred to nitrocellulose membranes. Total protein was visualized with Ponceau S and used as the loading control. Membranes were blocked in 5% milk in TBS-T and incubated overnight at 4 °C with primary antibodies against p53 (mouse liver and hepatocytes: Leica Biosystems, cat# NCL-p53-CM5p, RRID:AB_563933; IMR90: Santa Cruz Biotechnology, cat# sc-126, RRID:AB_628082), CDKN1A/p21 (Abcam, cat# ab188224), GADD45A (Cell Signaling Technology, cat# 4632T), PCNA (Cell Signaling Technology, cat# 13110), phospho-ATR (Ser428) (Cell Signaling Technology, cat# 2853), STAT1 (Cell Signaling Technology, cat# 9172S), phospho-STAT1 (Tyr701) (Cell Signaling Technology, cat# 9167) and IL8/CXCL8 (Abcam, cat# ab18672, RRID:AB_444617), each used at 1:1,000 (further details in Supplementary Table 4), followed by HRP-conjugated secondary antibodies: goat anti-rabbit IgG (H + L) (Millipore, cat# AP307P, RRID:AB_92641) or, for the mouse monoclonal p53 antibody used in the IMR90 experiments, goat anti-mouse IgG, IgM (H + L) (Thermo Fisher Scientific, cat# 31446, RRID:AB_228318), each used at 1:15,000. Signal was detected with SuperSignal West Pico PLUS (Thermo Fisher Scientific, cat# 34580) or, where greater sensitivity was required, SuperSignal West Femto (Thermo Fisher Scientific, cat# 34095) chemiluminescent substrate, and imaged on a Bio-Rad ChemiDoc. Blot images were assembled and aligned in Sciugo and band intensities were quantified in ImageJ (Fiji), normalizing each band to the corresponding Ponceau signal.

## Histology and pathological assessment

Formalin-fixed liver tissue was paraffin embedded and sectioned using standard approaches. Hematoxylin and eosin (H&E) staining was performed by the Sanford Burnham Prebys histology core using standard approaches. H&E images were scored for liver pathology, including foci of immune cell infiltration (hepatitis), by a board-certified veterinary pathologist (K.K., Histopathology Core, La Jolla Institute for Immunology) blinded to age and treatment.

## Oil Red O staining and quantification

Oil Red O staining was performed by the Sanford Burnham Prebys histology core using standard approaches. Stained sections were digitized on a Leica Biosystems Aperio ScanScope slide scanner at 20× objective magnification (0.50 µm/pixel), and quantification was performed on the 4×-downsampled pyramid level (2.01 µm/pixel) using Python 3.9 with OpenSlide^84^ and OpenCV. Oil Red O–positive signal was segmented in HSV color space using fixed thresholds (H 0–10 or 170–180, S ≥ 50, V ≥ 50; OpenCV 8-bit scale), followed by a 2 × 2 pixel morphological opening to suppress isolated pixel noise. Total tissue area was defined as all pixels with any RGB channel below 200, and hepatic lipid content is reported as the percentage of tissue area occupied by Oil Red O–positive signal. Segmentation parameters were identical for all specimens, and no manual region selection or per-slide threshold adjustment was applied.

## Published human dermal fibroblast data

Bulk RNA-seq data from primary human dermal fibroblasts of young (ages 23–33, n = 4) and aged (ages 64–72, n = 5) donors were reanalyzed from a previously published dataset (GEO accessions GSE307377 and GSE331495)^40,41^. Donor recruitment, consent and culture conditions are described in that study. Reanalysis was performed as described under Bulk RNA-seq analysis.

## IMR90 culture, irradiation and siRNA KD

IMR90 primary human fibroblasts (ATCC, CCL-186) were cultured at 37 °C, 5% CO_2_ and 3.5% O2 in DMEM (Gibco, 10313-121) supplemented with 10% FBS (Corning, 35-0-11-CV), 1% penicillin-streptomycin (Gibco, 15140-122) and 2 mM glutamine (Gibco, 25030-081). Cells were tested routinely for mycoplasma and were negative. Cells at 20–30% confluence were irradiated with 20 Gy X-rays using an RS-2000 X-ray irradiator (Rad Source Technologies) (day 0). Cells were split once they returned to confluence on day 3, transfected on day 4 and collected on day 5. Cells were transfected with 25 nM siRNA in 0.8% DharmaFECT reagent (Dharmacon) according to the manufacturer’s instructions, using two individual siRNAs targeting TP53 (si-p53; Dharmacon siGENOME, D-003329-08-0005 and D-003329-26-0005) or a non-targeting control siRNA (siNTC; Dharmacon siGENOME, D-001210-03-20). Cells were returned to regular medium 18–20 h after transfection. Proliferating, non-irradiated IMR90 cells were included as a control.

## EdU incorporation assay

Cells were plated on PhenoPlate 96-well microplates (PerkinElmer) and pulsed with 10 µM EdU (TCI, E1057) for 24 h, fixed in 10% formalin for 15 min at room temperature and permeabilized with three washes of TBS containing 0.5% Triton X-100. EdU was detected by click chemistry for 30 min at room temperature in the dark using 50 µM FAM-azide (Abcam, ab146477), 1 mM CuSO4 (Acros Organics, 422870050) and 100 mM sodium L-ascorbate (Chem-Impex, 01436) in TBS/0.5% Triton X-100, with ascorbate added last. Cells were washed three times in TBS/0.5% Triton X-100, once in PBS and counterstained with 4′,6-diamidino-2-phenylindole (DAPI), as described previously^65^. Plates were imaged on a Nikon Ti2 microscope with automated image capture. Images were analyzed in NIS-Elements AR v5.21.03 using dark-background subtraction, thresholding, size exclusion and automated partitioning to identify nuclei, and the fraction of EdU-positive nuclei was normalized to total cell number; n = 4 biological replicates.

## RT-qPCR

RNA from IMR90 cells was isolated either with RNA Miniprep kit (Zymo Research, cat# R2050) according to the manufacturer’s protocol. RNA was reverse transcribed with RevertAid Reverse Transcriptase (Thermo Fisher Scientific, cat# EP0441) and RiboLock RNase Inhibitor (Thermo Fisher Scientific, cat# EO0381) in 5× reaction buffer for RT (Thermo Fisher Scientific). Gene expression was quantified by a standard SYBR-based approach (PowerUp SYBR Green Master Mix, Applied Biosystems, cat# A25741) on a QuantStudio 6 Flex Real-Time PCR System in 384-well format (Applied Biosystems, cat# 4485691), using primers for TP53 and CDKN1A (sequences in Supplementary Table 5). Relative expression was calculated by the 2^-DDCt method normalized to the geometric mean of GAPDH and RPL13 as reference genes; n = 3 biological replicates.

## Bulk RNA-seq library preparation and sequencing

Total RNA was extracted from snap-frozen whole liver with the Direct-zol RNA Miniprep kit (Zymo Research) and from cultured fibroblasts with the Quick-RNA Miniprep kit (Zymo Research, cat# R1055), in both cases according to the manufacturer’s protocol. RNA integrity was verified on a TapeStation; all samples had RIN > 7. Poly(A) RNA isolation and library preparation were performed with the Watchmaker mRNA Library Prep Kit (Watchmaker Genomics, cat# 7BK0001) using xGEN Stubby adapters (IDT, cat# 336338420) and xGEN 10 nt UDIs (IDT, cat# 336338436). Libraries were sequenced on the Element Biosciences AVITI platform using the AVITI 2 × 75 High Output Cloudbreak Freestyle Kit (Element Biosciences, cat# 860-00015).

## Bulk RNA-seq analysis

Raw reads were quality-checked with FastQC v0.11.8 and, for experiments containing adapter sequence, trimmed with Trim Galore v0.4.4_dev, a wrapper for Cutadapt^85^. Reads were aligned to the mm10 reference genome with STAR v2.5.3a^86^ and converted to gene-level counts with the analyzeRepeats.pl script in HOMER^87^. Genes with at least 10 counts in at least 4 samples were retained, and one old sgRosa26 sample identified as a principal component analysis (PCA) outlier was excluded (n = 19). Gene counts were normalized and tested for differential expression with DESeq2 v1.48.2^88^ (R v4.5.0). Genes with adjusted P < 0.05 and |log2 fold change| > 0.5 were considered differentially expressed (DEGs). Within each cluster, genes were ordered by hierarchical clustering (Ward linkage, Euclidean distance). Over-representation analysis (ORA) of GO biological process terms for each cluster was performed with clusterProfiler v4.16.0^89^ (enrichGO), and Benjamini-Hochberg correction (adjusted P < 0.05). Redundant terms were collapsed with the clusterProfiler simplify function (semantic similarity cutoff 0.7). Gene set enrichment analysis (GSEA)^48^ was performed with fgsea v1.34.2^90^ against MSigDB GO Biological Process gene sets; the Whitfield cell-cycle gene set^47^ (Supplementary Table 1, Whitfield cell cycle genes) was used for cell-cycle heatmaps. Heatmaps show row Z-scores of log2(CPM + 1) values and were generated in Python with matplotlib v3.9.4^91^ and SciPy v1.13.1^92^.

## Single-nucleus RNA-seq analysis

Single-nucleus multiome was performed for 18 wild-type mouse samples, including young (∼4 months, n = 6), old (∼22 months, n = 6) and very old (∼30 months, n = 6). The sgRosa26- and sgTrp53-treated livers and the vehicle- and HDM201-treated livers were profiled and processed using the same pipeline. Raw sequencing reads were processed using cellranger-arc (v2.0.2, 10x Genomics). Ambient RNAs were removed using CellBender (v0.3.0)^93^. Cells marked as doublets by DoubletFinder (v2.0.4)^94^ were removed. Additional data processing was performed using Seurat (v5.2.1)^95^ and Signac (v1.12.0)^96^. Cells with >10% mitochondrial reads, <200 genes detected, <1,000 ATAC reads or ATAC enrichment at transcription start site <1 were discarded. The samples were merged, normalized using log normalization and integrated using Harmony^97^. After cell clustering by Seurat, initial cell type annotation for each cluster was performed based on canonical cell type marker genes for hepatocytes, cholangiocytes, endothelial cells, hepatic stellate cells (HSCs), myeloid cells, T cells and B cells. For each initial cell type, subclustering was performed and expression of cell type markers was examined. Subclusters expressing multiple cell type markers were considered potential doublets and removed. The p53 target module score was calculated with Seurat AddModuleScore using the Fischer p53 target gene list^43^ (Supplementary Table 1). Cluster 4 and cluster 5 module scores were calculated in the same way; for each cell type, the gene set was first restricted to genes detected in at least 5% of the reference-group cells of that cell type. Cell-type proportions were calculated per mouse as the percentage of total nuclei.

## Single-nucleus ATAC-seq analysis and p53 target gene network

snATAC-seq fragments were processed with cellranger-arc; nuclei were filtered on TSS enrichment > 1 and > 200 unique fragments. Peaks were called per cell type with MACS2 and a union peak set was generated. To define candidate p53 target genes, all peaks accessible in young or aged hepatocytes (detected in ≥1% of nuclei per age group) were scanned for the p53 binding motif (MA0106.3) from JASPAR2020^54^ using motifmatchr (MOODS algorithm; background nucleotide frequencies from the peak set, mm10) at a *P* < 5 × 10^−5^ threshold. Each motif-containing peak was assigned to its nearest annotated transcription start site (mm10), retaining peaks whose target gene was expressed in hepatocytes (mean normalized expression >= 0.1). This set was intersected with DEGs from bulk RNA-seq of aged sgTrp53 livers, retaining only peak-linked genes whose expression changed upon p53 loss, yielding 194 candidate p53-activated (downregulated in old sgTrp53) and 45 candidate p53-repressed (upregulated in young sgTrp53) genes. GO biological process over-representation of these gene sets was tested with clusterProfiler.

## MERFISH spatial transcriptomics (MERSCOPE)

Fresh liver tissue was embedded in OCT, frozen in isopentane cooled on dry ice and cryosectioned at 10 µm onto MERSCOPE slides (Vizgen)^44^. Sections were fixed, permeabilized and hybridized with a custom 300-gene MERFISH panel (Supplementary Table 1) comprising senescence, p53 target, inflammation and interferon-stimulated genes together with cell-type identification markers. Samples were processed on the MERSCOPE instrument according to the manufacturer’s instructions. Two cohorts were profiled: young, old and geriatric wild-type liver (n = 6, 6 and 6 mice) and aged sgRosa26- and sgTrp53-treated liver (n = 3 mice per group).

## MERFISH data analysis

Cell segmentation was performed with Cellpose^98^, and transcripts were assigned to cells to generate a cell-by-gene matrix. Cells were retained if they contained more than 100 transcripts and more than 10 unique genes; for Trp53 KO and Rosa26 KO samples, only the >10 unique genes criterion was applied. Counts were normalized with SCTransform^99^, integrated across samples with Harmony^97^ and clustered and visualized by uniform manifold approximation and projection (UMAP) in Seurat (v5.0.1.9001). Clusters were annotated using a panel of liver cell-type marker genes and by submitting the 20 most cluster-specific genes of each cluster to CellKB^100^. p53 target and cell-cycle module scores were computed with AddModuleScore restricted to panel genes. p53-high hepatocytes were defined as the top 5% of per-cell p53 target module score among hepatocytes. For neighborhood analysis, the 10 nearest neighbors of each p53-high hepatocyte (all cell types, by centroid distance) were identified within each liver section and the observed frequency of each cell type among these neighbors was compared to the expected frequency from random permutation of the p53-high label among hepatocytes of the same section (100 permutations per section); results are reported as the median per-mouse log2 observed/expected ratio. Infiltrated and non-infiltrated regions were selected manually in QuPath^101^ on the basis of small immune cell density; raw counts from cells within these regions were summed into pseudobulk profiles and normalized using the DESeq2^88^ median-of-ratios method.

## Tissue sectioning and slide preparation for CosMx

Paraffin-embedded liver blocks were fixed in 10% neutral buffered formalin or 4% paraformaldehyde and were less than 3 years old at the time of sectioning. The Sanford Burnham Prebys Histology Core cut 10 serial 5 µm sections from each block using diethyl pyrocarbonate (DEPC)-treated water, with all equipment and tools cleaned with RNaseZap. Sections were mounted in the center of Leica Bond Plus slides (Leica Biosystems, cat# S21.2113.A) and air-dried overnight at room temperature. Slides were stored at 4 °C with desiccant and shipped on ice packs to NanoString Technologies (Seattle, WA) for CosMx spatial molecular imaging. The second serial section was stained with hematoxylin and eosin, and a whole-slide image of it was used to select fields of view for profiling.

## CosMx data processing and analysis

Raw image segmentation and initial processing were performed by the Bruker team. Raw high-resolution CosMx images were segmented to infer cell boundaries. Transcripts were then assigned to cells accordingly to generate the initial cell-by-gene expression matrix.

CosMx data were subjected to stringent quality control. Data processing and downstream analyses were performed in Seurat (v5.1.0). Cells were retained if they contained at least 20 but fewer than 1,000 transcripts, at least 5 but fewer than 400 detected genes, and a cell area less than five times the geometric mean cell area.

Potential doublets arising from segmentation of two or more cells as a single cell were identified with Scrublet (v0.2.3)^102^, which detects doublets from gene expression profiles. Scrublet was applied independently to each field of view (FOV). On average, Scrublet identified around 5 potential doublets per FOV, corresponding to 0.3% cells on average per FOV. Given the low predicted doublet rate, no cells were removed on the basis of Scrublet predictions, to avoid excluding true singlets through false-positive doublet calls.

Cells that passed quality control (QC) were further normalized, integrated, clustered and annotated. The normalization was performed using SCT transform^99^ in Seurat^103^ with “vst.flavor=v2” and using “vars.to.regress” to remove confounding factors including number of genes and number of transcripts. Different samples and replicates were integrated using “IntegrateData” in Seurat after identifying a subset of anchors with “FindIntegrationAnchors”. PCA was performed on the integrated data. Based on variance explained by each principal component (PC), we selected the top 20 PCs for generating the UMAP^104^ embedding, building neighborhood graphs and also cell clustering using “FindClusters” with “resolution=0.2”. Cell type annotation was performed and validated by marker gene expression, top DEGs of each cluster identified using Wilcoxon rank-sum test with Bonferroni correction, and cell marker protein staining signals.

## Cell–cell interaction analysis of CosMx data

Spatially proximal cell–cell interactions were inferred from the CosMx data using CellChat (v2.1.2)^105^ with the curated mouse CellChatDB interaction database^51^. Communication probabilities were calculated with the "computeCommunProb" function using default parameters except type = "truncatedMean", interaction.range = 250, contact.range = 10 and scale.distance = 3.29, as suggested by CellChat during an initial test run. Interactions involving cell types represented by fewer than 10 cells were excluded with the "filterCommunication" function. Inferred communication patterns were compared between groups and visualized with the "netVisual_bubble" function.

## Statistics and reproducibility

No statistical method was used to predetermine sample size; sample sizes were based on previous studies in the lab. Statistical tests, multiple-comparison corrections and n (biological replicates) for each experiment are stated in the figure legends; the longitudinal frailty analysis is described in Statistical analysis of frailty. Analyses were performed in GraphPad Prism (v10.4.2) and R (v4.5.0).

## Statistical analysis of frailty

Longitudinal changes in frailty index were analyzed with a linear mixed model for repeated measures, fitted by restricted maximum likelihood (REML) with the nlme package (v3.1-168; doi:10.32614/CRAN.package.nlme) in R (v4.5.0). Group (young vehicle, old vehicle, old HDM201), timepoint (baseline and weeks 5, 10, 15 and 19, as a categorical variable) and their interaction were included as fixed effects. Repeated measurements within each mouse were modeled with an unstructured covariance matrix, allowing a separate variance for each timepoint and group and a separate correlation for each pair of timepoints. This structure fitted the data better than models with a random intercept, or a random intercept and slope, per mouse (lowest Akaike information criterion), and it does not assume equal variance across timepoints (sphericity). All mice were included. Mice that died or were euthanized during the study contributed all measurements obtained before death; the model uses all available observations without imputation. Within each group, frailty at each post-baseline timepoint was compared with baseline using linear contrasts of the model estimates (t-tests), with Šídák correction for the four comparisons per group.

## Use of AI tools

During preparation of this manuscript, the authors used Claude (Anthropic) models Opus 4.8, Opus 5 and Opus 5.5 to edit writing, assist with literature search and code writing. No AI tool was used to directly generate data, figures or scientific conclusions. The authors reviewed and edited all AI-assisted output and take full responsibility for the content of the manuscript.

## Data availability

Bulk RNA-seq data from young and old sgRosa26- and sgTrp53-treated mouse livers on normal diet and from aged sgRosa26- and sgTrp53-treated livers on high-fat diet have been deposited in the Gene Expression Omnibus (GEO) under accessions GSE314411 and GSE314410, respectively. Bulk RNA-seq data from irradiated IMR90 fibroblasts transfected with control or TP53 siRNA are available in GEO under accession GSE353900. MERFISH data from aged sgRosa26- and sgTrp53-treated livers and CosMx spatial transcriptomics data from young and aged sgRosa26- and sgTrp53-treated livers are available in GEO under accessions GSE339101 and GSE315907, respectively. 10X single-nucleus multiome (snRNA-seq and snATAC-seq) data from young, old and geriatric wild-type livers, young and old sgTrp53-treated livers and aged vehicle- and HDM201-treated livers are available through the SenNet Data Portal under dataset IDs SNT296.PSCN.662 (female) and SNT987.ZGSS.452 (male; https://doi.org/10.60586/SNT987.ZGSS.452), SNT757.ZBHZ.862 and SNT638.DJNJ.996, respectively, where the full nuclei isolation, library preparation and sequencing protocols for these datasets are also deposited. MERFISH data from young, old and geriatric wild-type livers are available from the corresponding author upon request. Previously published bulk RNA-seq data from primary human dermal fibroblasts of young and aged donors that were reanalyzed in this study are available in GEO under accession GSE307377; the same samples are also deposited within GSE331495. A summary of all datasets generated and reanalyzed in this study, with their repositories and accession numbers, is provided in Supplementary Table 6.

## Supporting information

Supplementary Table 1: Gene lists (p53 target genes, ISGs, Whitfield cell-cycle genes, SenNet MERFISH panel, p53 KO old MERSCOPE panel)

Supplementary Table 2: DEG clusters and GO biological process terms, p53 knockout mouse liver (Fig. 3a)

Supplementary Table 3: DEG clusters and GO biological process terms, TP53 knockdown in irradiated IMR90 cells (Extended Fig. 4e)

Supplementary Table 4: Antibodies

Supplementary Table 5: qPCR primers

## Acknowledgements

We thank Adriana Charbono and Buddy Charbono for technical support with mouse experiments, and Kenneth Kim, DVM, Dipl. ACVP (Director, Histopathology Core, La Jolla Institute for Immunology) for histopathological assessment of liver sections. We thank the SBP Bioinformatics Core (Rabi Murad and Neelam Sinha) and the SBP Functional Genomics Core (Chun-Teng Huang) for technical assistance. The research performed by the SBP Functional Genomics Core in this publication was supported by the Shared Instrumentation Grant S10 OD036254. All SBP NCI-designated Cancer Center core facilities were supported by the NCI Cancer Center Support Grant P30 CA030199.

Work in the laboratory of P.D.A. was supported by P01 AG031862, P01 AG073084 and the SenNet grant U54 AG079758. R.A., M.G.T. and G.G. were supported by the California Institute for Regenerative Medicine grant EDUC4-12813. M.G.T. was also supported by an American Society of Hematology (ASH) Hematology Inclusion Pathway (HIP) Graduate Student Award. K.M. was supported by NIH K99 AG073450. A.S. was supported by NIH R01 AR077448. A.R.-A. was supported by a donation from Mr. Javier Moreno (II040141) and a mobility grant from the Spanish RIS Association. RNA-seq of primary human dermal fibroblasts was performed through the San Diego Nathan Shock Center, supported by NIH P30 AG068635. We thank all members of the Adams lab for critical discussions.

## Author contributions

M.G.T. and K.M. conceived and designed the study, performed experiments, analyzed and interpreted data, and wrote the manuscript. R.A., A.H., A.R., A.G., J.P., G.G., L.H., I.H., M.A., A.D., A.R.-A., C.M., C.C.M., E.H., S.P., M.S., G.Ga., H.T., T.M. and Ca.K. performed mouse experiments and provided reagents, resources or expertise. K.Y.L., K.L., S.M., C.H.S., G.E., X.L., J.C., Co.K. and J.O. performed computational analyses. Ap.W., B.R., Al.W., N.Z., Q.Z., K.Y.Y., R.A.P., A.S., K.M.T. and Y.X.W. supervised work in their respective laboratories. E.S. contributed to project coordination. P.D.A. conceived and supervised the study, acquired funding and edited the manuscript. All authors reviewed and approved the final manuscript.

## Competing interests

The authors declare no competing interests.

## Abbreviations

AAV: adeno-associated virus
ALT: alanine aminotransferase
ANOVA: analysis of variance
AP: antagonistic pleiotropy
ATAC-seq: assay for transposase-accessible chromatin using sequencing
BH: Benjamini–Hochberg
BP: biological process
CBC: complete blood count
CKI: cyclin-dependent kinase inhibitor
CPM: counts per million
CRISPR: clustered regularly interspaced short palindromic repeats
DAPI: 4′,6-diamidino-2-phenylindole
DEG: differentially expressed gene
DEPC: diethyl pyrocarbonate
DMEM: Dulbecco’s modified Eagle medium
EdU: 5-ethynyl-2′-deoxyuridine
EMM: estimated marginal mean
FBS: fetal bovine serum
FOV: field of view
GEO: Gene Expression Omnibus
GO: Gene Ontology
GSEA: gene set enrichment analysis
H&E: hematoxylin and eosin
HBSS: Hanks’ balanced salt solution
HFD: high-fat diet
HGB: hemoglobin
HSC: hepatic stellate cell
ICE: Inference of CRISPR Edits
IQR: interquartile range
IR: irradiated
ISG: interferon-stimulated gene
KO: knockout
L-R: ligand–receptor
LME: linear mixed-effects
LYM: lymphocytes
MASH: metabolic dysfunction-associated steatohepatitis
MASLD: metabolic dysfunction-associated steatotic liver disease
Mdm2i: MDM2 inhibitor
MERFISH: multiplexed error-robust fluorescence in situ hybridization
MHC-II: major histocompatibility complex class II
MNAR: missing not at random
MON: monocytes
MSigDB: Molecular Signatures Database
NAFLD: nonalcoholic fatty liver disease
NASH: nonalcoholic steatohepatitis
NES: normalized enrichment score
NEU: neutrophils
NTC: non-targeting control
OCT: optimal cutting temperature compound
ORA: over-representation analysis
PAM: protospacer adjacent motif
PBS: phosphate-buffered saline
PC: principal component
PCA: principal component analysis
PCNA: proliferating cell nuclear antigen
PLT: platelets
QC: quality control
RBC: red blood cells
RT-qPCR: reverse transcription quantitative
PCR saCas9: *Staphylococcus aureus* Cas9
SASP: senescence-associated secretory phenotype
SCT: sctransform normalization
SD: standard deviation
SEM: standard error of the mean
sgRNA: single-guide RNA
siNTC: non-targeting control siRNA
snATAC-seq: single-nucleus ATAC-seq
snRNA-seq: single-nucleus RNA sequencing
TBG: thyroxine-binding globulin
TBS: Tris-buffered saline
TSS: transcription start site
UMAP: uniform manifold approximation and projection
WBC: white blood cells

---

**Extended Figure 1:**
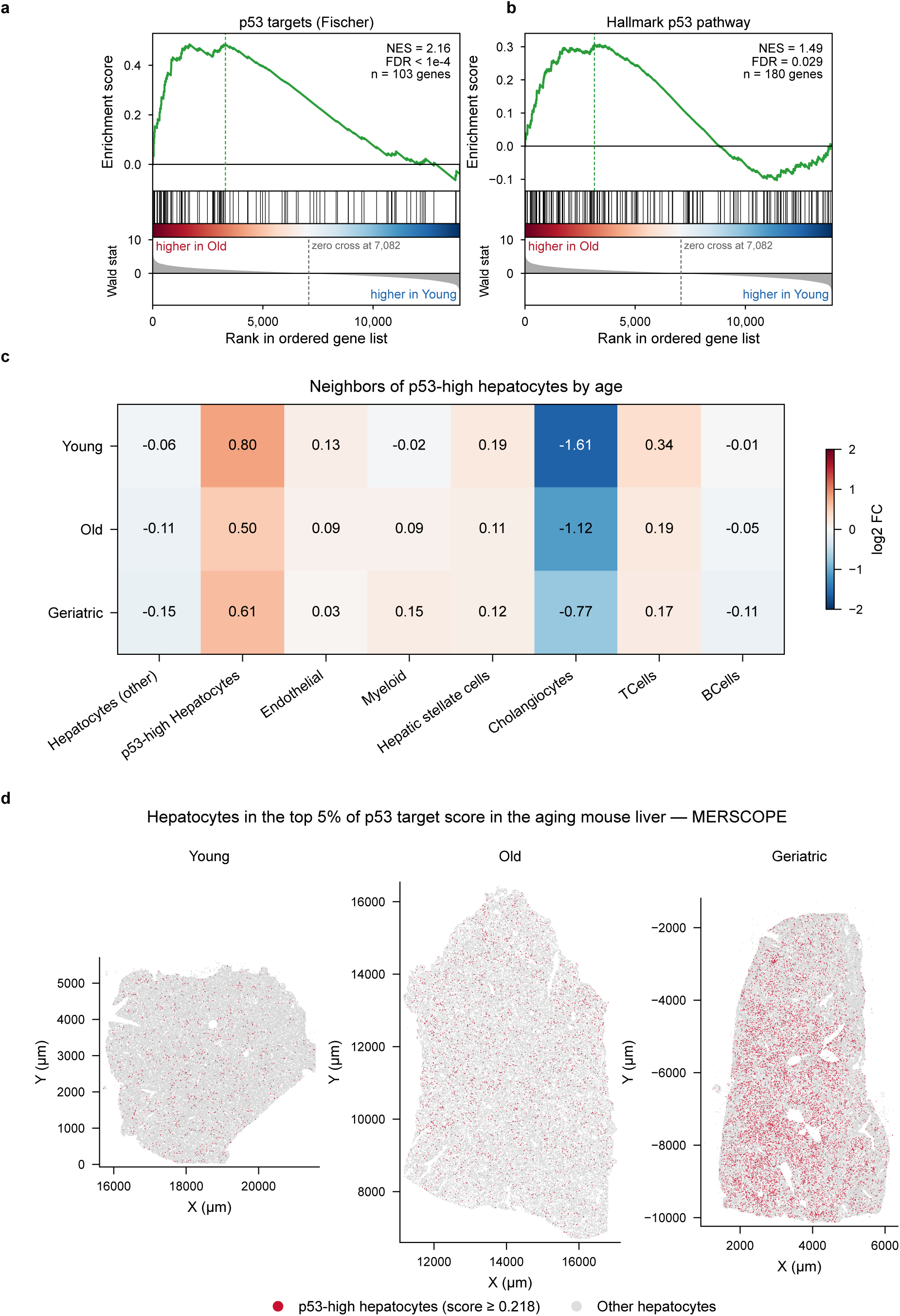
p53 activity increases with age in human fibroblasts and mouse hepatocytes. a, b, GSEA of p53 target genes in aged (ages 64–72, n = 5 donors) compared with young (ages 23–33, n = 4 donors) primary human dermal fibroblasts. Fischer p53 target genes (a) and MSigDB Hallmark p53 pathway (b). **c,** Cell-type composition of the neighborhood of p53-high hepatocytes by age, calculated from the MERFISH dataset (Young, n = 4 mice; Old, n = 4 mice; Geriatric, n = 6 mice). p53-high hepatocytes are defined as the top 5% of per-cell p53 target module score in hepatocytes. Values show the median per-mouse log2 fold change of observed over expected neighbor frequency. Red indicates enrichment and blue indicates depletion. **d,** Spatial distribution of p53-high hepatocytes in representative liver sections from young, old and geriatric mice from the MERFISH dataset. p53-high hepatocytes (red) are defined as in c. All other hepatocytes are shown in grey. Each section displays the same number of randomly sampled hepatocytes (n = 72,601).

**Extended Figure 2:**
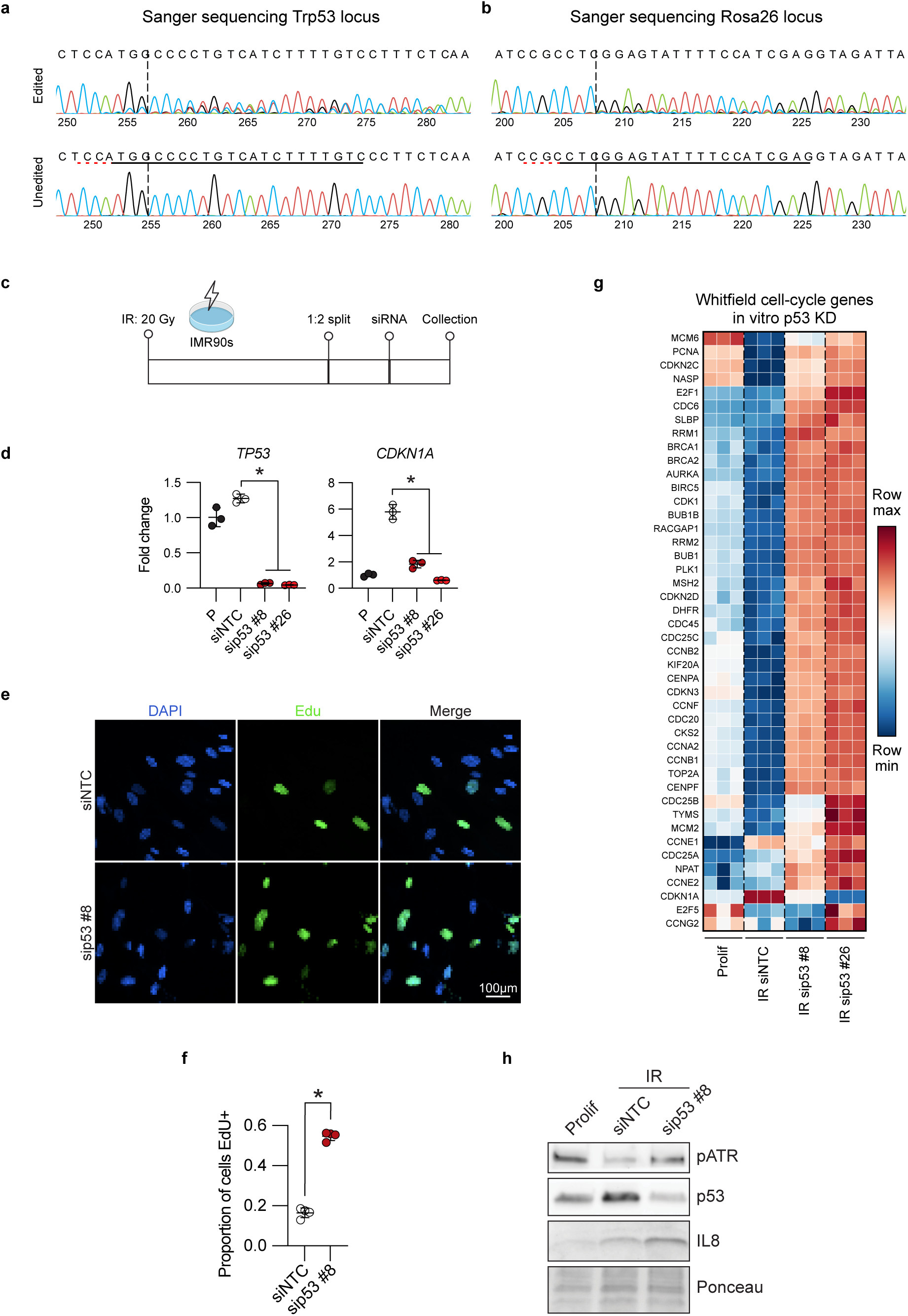
Validation of in vivo sgRNAs and of TP53 KD in irradiated IMR90 cells. a, b, Representative Sanger sequencing chromatograms of the loci targeted by sgTrp53 (a) and sgRosa26 (b) in genomic DNA from whole liver. Top, edited locus; bottom, corresponding control. sgRNA target sequence, underlined; PAM, horizontal dashed line; predicted cut site, vertical dashed line. **c,** Schematic representation of the in vitro experimental design. IMR90 fibroblasts were irradiated with 20 Gy on day 0, split 1:2 on day 3, transfected with siRNA on day 4 and collected on day 5. **d,** qPCR of TP53 and its target gene CDKN1A after siRNA KD, n = 3. **e,** Representative images of the EdU assay after TP53 KD. **f,** Quantification of the EdU assay normalized to cell number, n = 4. g, Heatmap of Whitfield proliferation genes by bulk RNA-seq, n = 3. **h,** Western blot of p-ATR, p53 and IL8, n = 1. Ponceau was used as loading control. (d, f) Statistical significance was assessed with two-tailed Student’s t-tests; each dot is a biological replicate. Data shown as means ±SD. \**P* < 0.05. Prolif: proliferating control; NTC: non-targeting control.

**Extended Figure 3:**
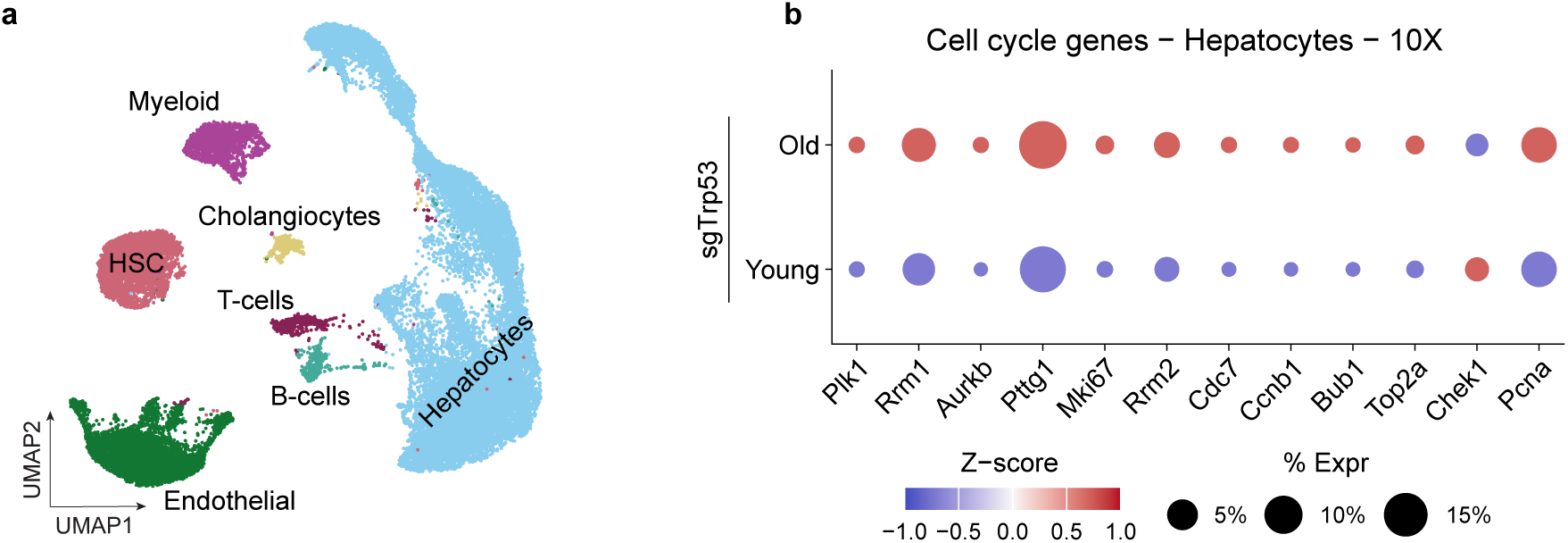
Single-nucleus analysis of young and aged sgTrp53-treated livers. **a**, UMAP representing 10X multiome profiling of young and old sgTrp53-treated livers, n = 2 mice per age group. **b,** Dot plot of cell-cycle genes in young and aged sgTrp53 hepatocytes (Young sgTrp53, n = 10,589 cells from 2 mice; Old sgTrp53, n = 7,664 cells from 2 mice).

**Extended Figure 4:**
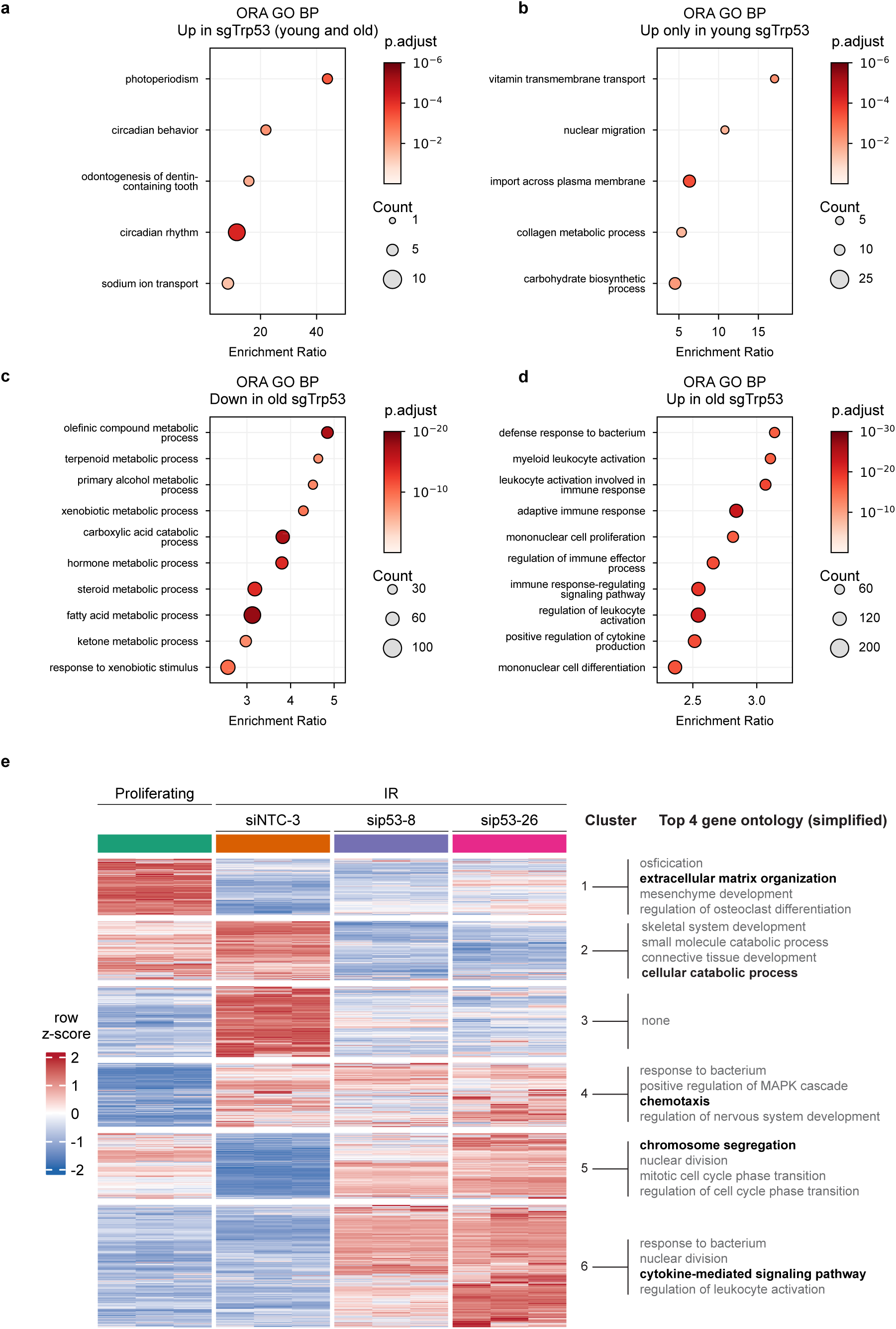
Pathway analysis of the effects of p53 loss in vivo and effect of TP53 KD in irradiated IMR90 cells. a–d, Over-representation analysis (ORA) of GO BP terms among genes differentially expressed in sgTrp53 versus sgRosa26 found by bulk RNA-seq. Genes up in both young and old sgTrp53 (a), up only in young sgTrp53 (b), down in old sgTrp53 (c) and up in old sgTrp53 (d). Position on the x-axis indicates the enrichment ratio; dot size indicates the number of DEGs per term; color indicates the adjusted *P* value. **e,** Heatmap of DEGs after TP53 KD in irradiated IMR90 cells by bulk RNA-seq, n = 3 biological replicates per group. The four most significant simplified GO BP terms are shown per cluster. Cluster 3 returned no significantly enriched term. IR, irradiated; NTC, non-targeting control.

**Extended Figure 5:**
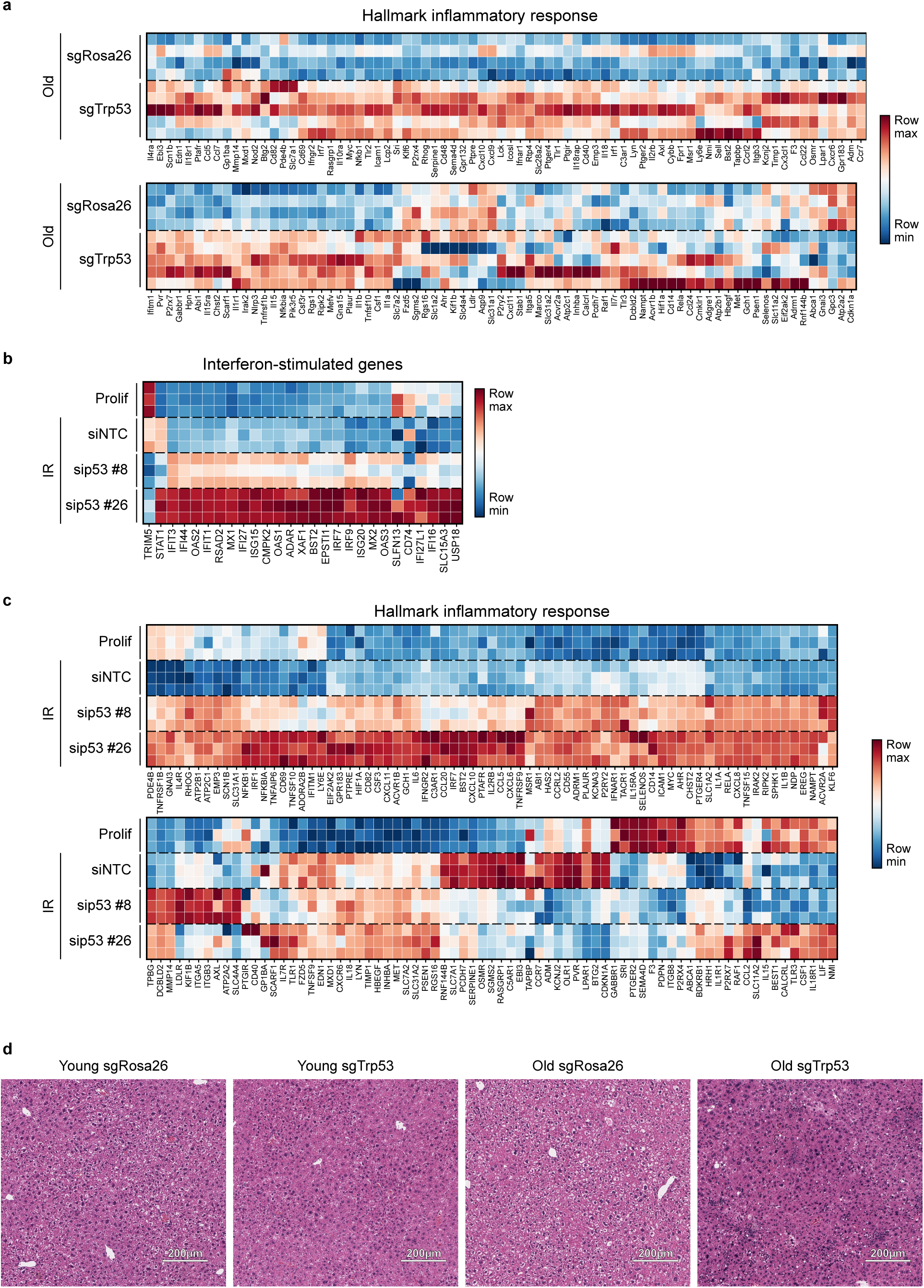
Inflammatory profiling after modulation of p53. a, Heatmap of Hallmark inflammatory response genes in old sgRosa26-vs old sgTrp53-treated livers (Old sgRosa26, n = 4; Old sgTrp53, n = 5). **b,** Heatmap of ISGs in irradiated IMR90 cells after TP53 KD. **c,** Heatmap of Hallmark inflammatory response genes in irradiated IMR90 cells after TP53 KD. (b, c) n = 3 biological replicates per group. **d,** Representative images of H&E staining in young and aged sgRosa26-or sgTrp53-treated livers. Scale bar, 200 µm.

**Extended Figure 6:**
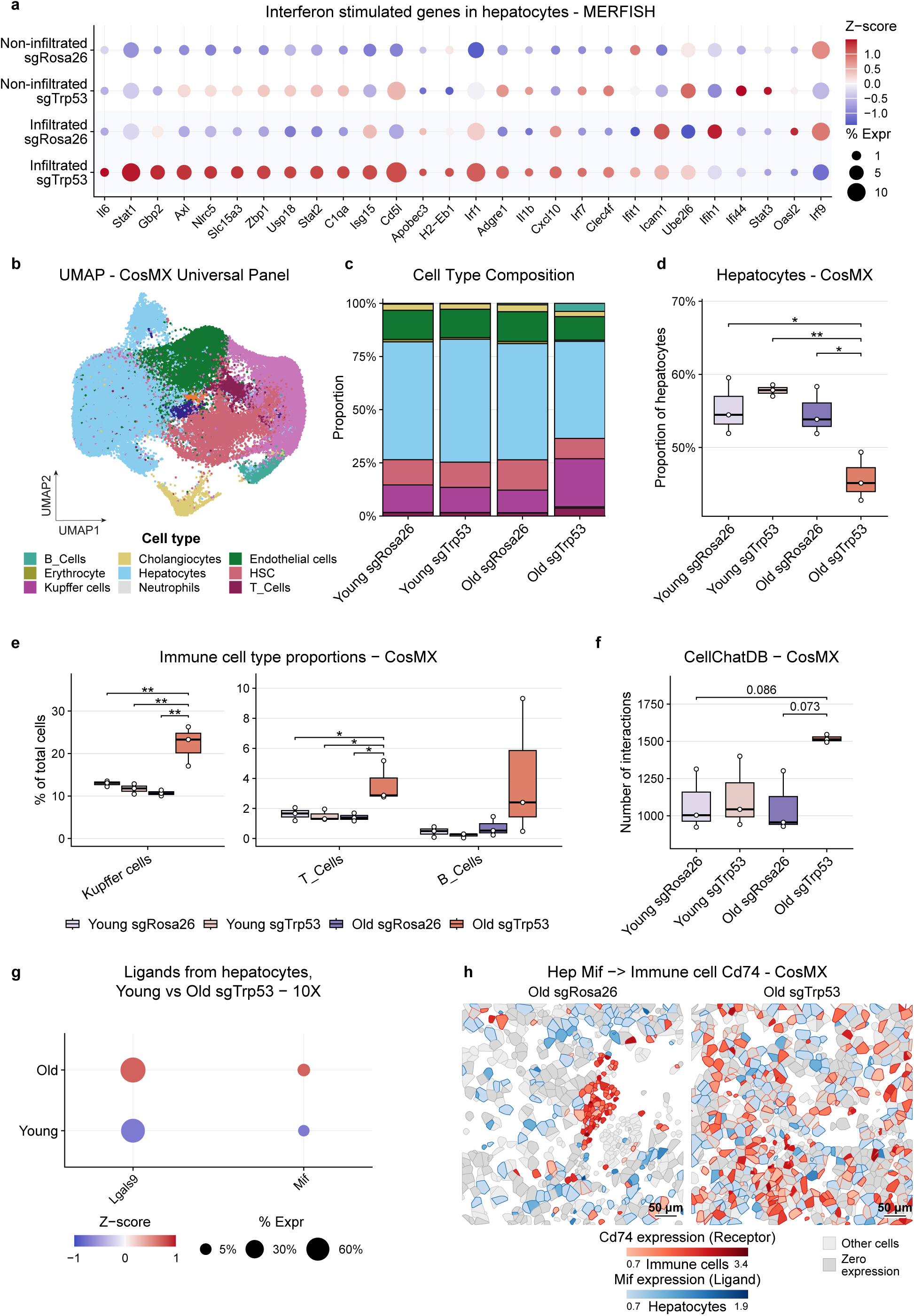
Single-cell spatial profiling of immune infiltration in aged sgTrp53 livers. a, Dot plot of ISGs in hepatocytes from non-infiltrated and immune-infiltrated regions of aged sgRosa26 and sgTrp53-treated livers, profiled by MERFISH. n = 3 mice per sgRNA treatment; non-infiltrated sgRosa26, n = 15,091 cells; non-infiltrated sgTrp53, n = 7,798 cells; infiltrated sgRosa26, n = 3,563 cells; infiltrated sgTrp53, n = 4,519 cells. **b,** UMAP representing CosMx spatial transcriptomics profiling of young and aged sgRosa26-and sgTrp53-treated livers with the Mouse Universal Cell Characterization RNA Panel. **c,** Bar plot showing the cell-type composition of young and aged sgRosa26-and sgTrp53-treated livers from the CosMx dataset. (b, c) n = 3 mice per age group. **d,** Box plot showing the proportion of hepatocytes in the CosMx dataset. *P* values: young sgRosa26 vs old sgTrp53, *P* = 0.021; young sgTrp53 vs old sgTrp53, *P* = 0.0058; old sgRosa26 vs old sgTrp53, *P* = 0.030. **e,** Box plots showing the proportions of Kupffer cells, T cells and B cells, as a percentage of all cells, in the CosMx dataset. *P* values, Kupffer cells: young sgRosa26 vs old sgTrp53, *P* = 0.0077; young sgTrp53 vs old sgTrp53, *P* = 0.0036; old sgRosa26 vs old sgTrp53, *P* = 0.0020; T cells: young sgRosa26 vs old sgTrp53, *P* = 0.049; young sgTrp53 vs old sgTrp53, *P* = 0.037; old sgRosa26 vs old sgTrp53, *P* = 0.029; no significant differences in B cells. **f,** Box plot showing the number of significant ligand–receptor interactions inferred from CellChatDB in the CosMx dataset. *P* values: young sgRosa26 vs old sgTrp53, *P* = 0.086; old sgRosa26 vs old sgTrp53, *P* = 0.073. **g,** Dot plot of *Lgals9* and *Mif* in young and aged sgTrp53 hepatocytes from the 10X multiome dataset (Young sgTrp53, n = 10,589 cells from 2 mice; Old sgTrp53, n = 7,664 cells from 2 mice). **h,** CosMx cell segmentation maps of *Cd74* in immune cells (red gradient) and *Mif* in hepatocytes (blue gradient). Other cell types are shown as light grey and zero-expressing cells as medium grey. Scale bar, 50 µm. (d–f) Each dot is one biological replicate (n = 3 mice per group). Box shows the 25th–75th percentile; the center line shows the median; whiskers extend to 1.5 × IQR. (d–f) Statistical significance was assessed with one-way ANOVA with Tukey’s correction for multiple comparisons. \**P* < 0.05, \*\**P* < 0.01.

**Extended Figure 7:**
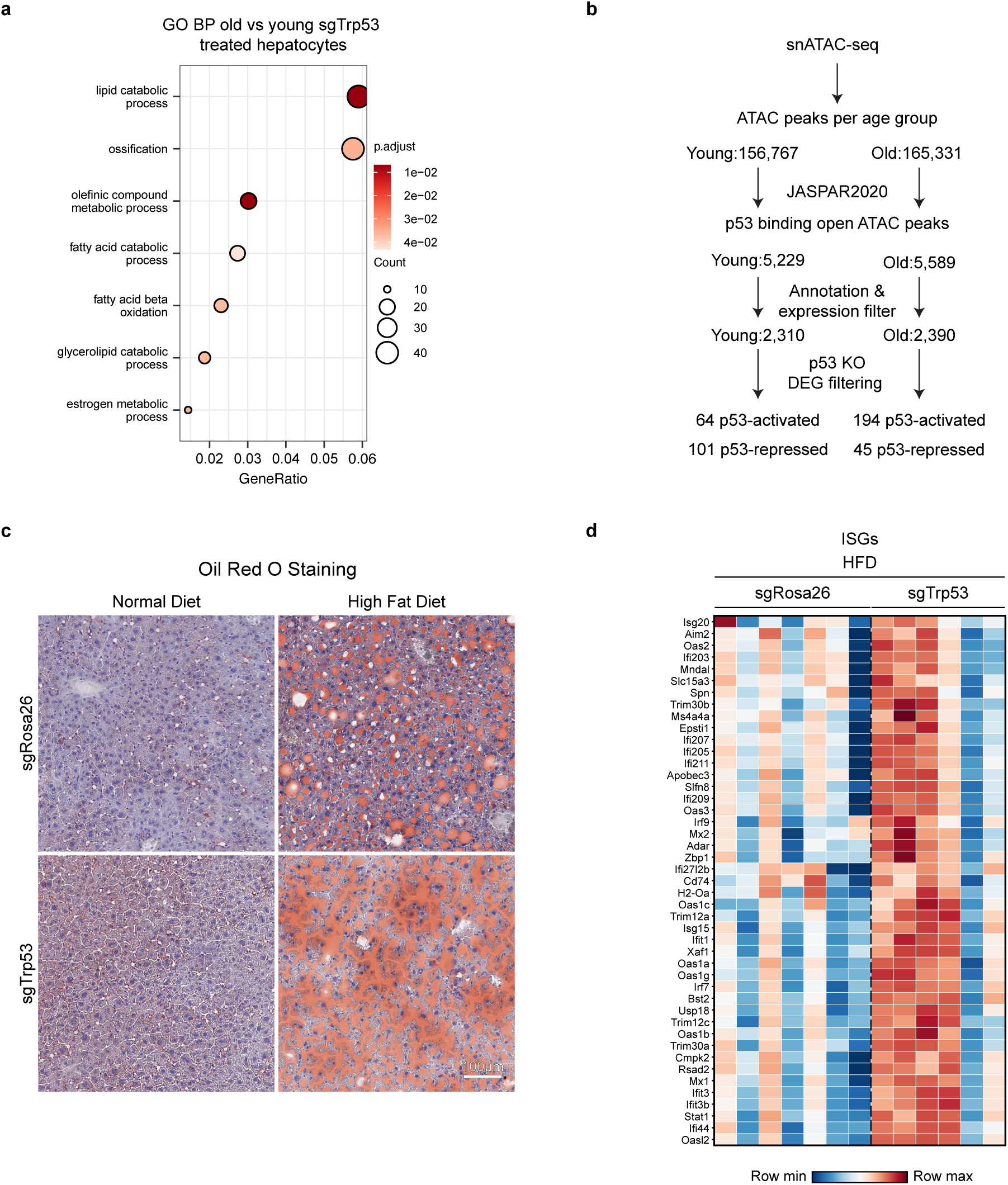
p53 controls the catabolic program of old hepatocytes and inhibits interferon signaling on a high-fat diet. a, ORA GO BP of genes downregulated in old vs young sgTrp53 hepatocytes. Terms with adjusted *P* < 0.05 are shown, ordered by gene ratio. **b,** Schematic representation of the workflow used to define candidate p53 target genes in young and aged hepatocytes. **c,** Representative Oil Red O-stained liver sections from aged mice treated with sgRosa26 or sgTrp53 fed either normal diet or HFD. Scale bar, 100 µm. **d,** Heatmap of ISGs from bulk RNA-seq of aged sgRosa26 and sgTrp53-treated livers on HFD (Old HFD sgRosa26, n = 7; Old HFD sgTrp53, n = 6).

**Extended Figure 8:**
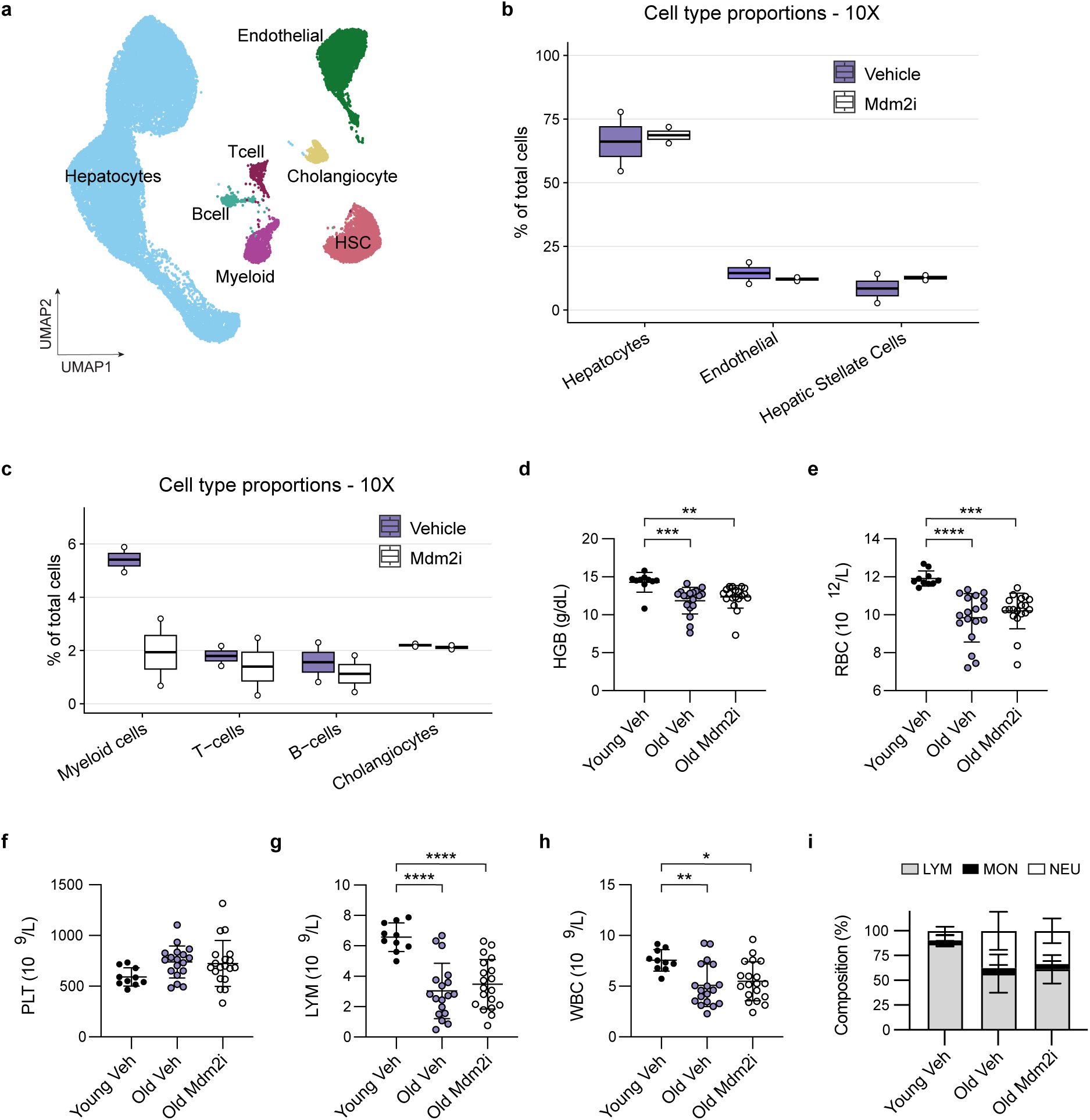
MDM2 inhibition does not cause hematological toxicity in aged mice. a, UMAP representing 10X multiome profiling of old female mice treated with HDM201 (Mdm2i) or vehicle, annotated by cell type (Old Veh, n = 17,247 cells from 2 mice; Old Mdm2i, n = 17,205 cells from 2 mice). **b, c,** Box plots showing the proportions of hepatocytes, endothelial cells and HSCs (b) and of myeloid cells, T cells, B cells and cholangiocytes (c) in vehicle-vs Mdm2i-treated aged livers from the 10X multiome dataset. **d–i,** CBC after 16 weeks of treatment in young vehicle-, old vehicle-and old Mdm2i-treated mice (Young Veh, n = 10; Old Veh, n = 18; Old Mdm2i, n = 19): hemoglobin (HGB; d), red blood cells (RBC; e), platelets (PLT; f), lymphocytes (LYM; g) and white blood cells (WBC; h). **d,** HGB, *P* values: Young Veh vs Old Veh, *P* = 0.0009; Young Veh vs Old Mdm2i, *P* = 0.0080. **e,** RBC, *P* values: Young Veh vs Old Veh, *P* < 0.0001; Young Veh vs Old Mdm2i, *P* = 0.0002. **g,** LYM, *P* values: Young Veh vs Old Veh, *P* < 0.0001; Young Veh vs Old Mdm2i, *P* < 0.0001. **h,** WBC, *P* values: Young Veh vs Old Veh, *P* = 0.0046; Young Veh vs Old Mdm2i, *P* = 0.0180. **i,** White blood cell differential, shown as the percentage of LYM, monocytes (MON) and neutrophils (NEU). (b, c) Box shows the 25th–75th percentile; the center line shows the median; whiskers extend to 1.5 × IQR; no statistical test was performed owing to n = 2 mice per group. (d–h) Statistical significance was assessed with one-way ANOVA with Tukey’s correction for multiple comparisons. (d–i) Data shown as means ±SD. \**P* < 0.05, \*\**P* < 0.01, \*\*\**P* < 0.001, \*\*\*\**P* < 0.0001.

## Notes

### Competing Interest Statement

The authors have declared no competing interest.

## References

1 Lane, D. P. Cancer. p53, guardian of the genome. Nature 358, 15–16 (1992). 10.1038/358015a0

2 el-Deiry, W. S., et al. WAF1, a potential mediator of p53 tumor suppression. Cell 75, 817–825 (1993). 10.1016/0092-8674(93)90500-p

3 Harper, J. W., Adami, G. R., Wei, N., Keyomarsi, K. & Elledge, S. J. The p21 Cdk-interacting protein Cip1 is a potent inhibitor of G1 cyclin-dependent kinases. Cell 75, 805–816 (1993). 10.1016/0092-8674(93)90499-g

4 Xiong, Y. et al. p21 is a universal inhibitor of cyclin kinases. Nature 366, 701–704 (1993). 10.1038/366701a0

5 Gu, Y., Turck, C. W. & Morgan, D. O. Inhibition of CDK2 activity in vivo by an associated 20K regulatory subunit. Nature 366, 707–710 (1993). 10.1038/366707a0

6 Yonish-Rouach, E. et al. Wild-type p53 induces apoptosis of myeloid leukaemic cells that is inhibited by interleukin-6. Nature 352, 345–347 (1991). 10.1038/352345a0

7 Serrano, M., Lin, A. W., McCurrach, M. E., Beach, D. & Lowe, S. W. Oncogenic ras provokes premature cell senescence associated with accumulation of p53 and p16INK4a. Cell 88, 593–602 (1997). 10.1016/s0092-8674(00)81902-9

8 Sugrue, M. M., Shin, D. Y., Lee, S. W. & Aaronson, S. A. Wild-type p53 triggers a rapid senescence program in human tumor cells lacking functional p53. Proc Natl Acad Sci U S A 94, 9648–9653 (1997). 10.1073/pnas.94.18.9648

9 Kastenhuber, E. R. & Lowe, S. W. Putting p53 in context. Cell 170, 1062–1078 (2017). 10.1016/j.cell.2017.08.028

10 Kandoth, C. et al. Mutational landscape and significance across 12 major cancer types. Nature 502, 333–339 (2013). 10.1038/nature12634

11 Kruiswijk, F., Labuschagne, C. F. & Vousden, K. H. p53 in survival, death and metabolic health: a lifeguard with a licence to kill. Nat Rev Mol Cell Biol 16, 393–405 (2015). 10.1038/nrm4007

12 López-Otín, C., Blasco, M. A., Partridge, L., Serrano, M. & Kroemer, G. Hallmarks of aging: An expanding universe. Cell 186, 243–278 (2023). 10.1016/j.cell.2022.11.001

13 Tyner, S. D. et al. p53 mutant mice that display early ageing-associated phenotypes. Nature 415, 45–53 (2002). 10.1038/415045a

14 Edwards, M. G. et al. Gene expression profiling of aging reveals activation of a p53-mediated transcriptional program. BMC Genomics 8, 80 (2007). 10.1186/1471-2164-8-80

15 Tyshkovskiy, A. et al. Universal transcriptomic hallmarks of mammalian ageing and mortality. Nature 654, 173–188 (2026). 10.1038/s41586-026-10542-3

16 Meletis, K. et al. p53 suppresses the self-renewal of adult neural stem cells. Development 133, 363–369 (2006). 10.1242/dev.02208

17 Gil-Perotin, S. et al. Loss of p53 induces changes in the behavior of subventricular zone cells: implication for the genesis of glial tumors. J Neurosci 26, 1107–1116 (2006). 10.1523/JNEUROSCI.3970-05.2006

18 Liu, Y. et al. p53 regulates hematopoietic stem cell quiescence. Cell Stem Cell 4, 37–48 (2009). 10.1016/j.stem.2008.11.006

19 Xue, W. et al. Senescence and tumour clearance is triggered by p53 restoration in murine liver carcinomas. Nature 445, 656–660 (2007). 10.1038/nature05529

20 Ventura, A. et al. Restoration of p53 function leads to tumour regression in vivo. Nature 445, 661–665 (2007). 10.1038/nature05541

21 Coppé, J. P. et al. Senescence-associated secretory phenotypes reveal cell-nonautonomous functions of oncogenic RAS and the p53 tumor suppressor. PLoS Biol 6, 2853–2868 (2008). 10.1371/journal.pbio.0060301

22 Acosta, J. C. et al. Chemokine signaling via the CXCR2 receptor reinforces senescence. Cell 133, 1006–1018 (2008). 10.1016/j.cell.2008.03.038

23 Kuilman, T. et al. Oncogene-induced senescence relayed by an interleukin-dependent inflammatory network. Cell 133, 1019–1031 (2008). 10.1016/j.cell.2008.03.039

24 Franceschi, C. et al. Inflamm-aging. An evolutionary perspective on immunosenescence. Ann N Y Acad Sci 908, 244–254 (2000). 10.1111/j.1749-6632.2000.tb06651.x

25 De Cecco, M. et al. L1 drives IFN in senescent cells and promotes age-associated inflammation. Nature 566, 73–78 (2019). 10.1038/s41586-018-0784-9

26 Williams, G. C. Pleiotropy, natural selection, and the evolution of senescence. Evolution 11, 398–411 (1957). 10.1111/j.1558-5646.1957.tb02911.x

27 Maier, B. et al. Modulation of mammalian life span by the short isoform of p53. Genes Dev 18, 306–319 (2004). 10.1101/gad.1162404

28 Dimri, G. P. et al. A biomarker that identifies senescent human cells in culture and in aging skin in vivo. Proc Natl Acad Sci U S A 92, 9363–9367 (1995). 10.1073/pnas.92.20.9363

29 Baker, D. J. et al. Clearance of p16Ink4a-positive senescent cells delays ageing-associated disorders. Nature 479, 232–236 (2011). 10.1038/nature10600

30 Baker, D. J. et al. Naturally occurring p16(Ink4a)-positive cells shorten healthy lifespan. Nature 530, 184–189 (2016). 10.1038/nature16932

31 Xu, M. et al. Senolytics improve physical function and increase lifespan in old age. Nat Med 24, 1246–1256 (2018). 10.1038/s41591-018-0092-9

32 Beauséjour, C. M. et al. Reversal of human cellular senescence: roles of the p53 and p16 pathways. EMBO J 22, 4212–4222 (2003). 10.1093/emboj/cdg417

33 Wiley, C. D. et al. Small-molecule MDM2 antagonists attenuate the senescence-associated secretory phenotype. Sci Rep 8, 2410 (2018). 10.1038/s41598-018-20000-4

34 Miller, K. N. et al. p53 enhances DNA repair and suppresses cytoplasmic chromatin fragments and inflammation in senescent cells. Nat Commun 16, 2229 (2025). 10.1038/s41467-025-57229-3

35 García-Cao, I. et al. "Super p53" mice exhibit enhanced DNA damage response, are tumor resistant and age normally. EMBO J 21, 6225–6235 (2002). 10.1093/emboj/cdf595

36 Mendrysa, S. M. et al. Tumor suppression and normal aging in mice with constitutively high p53 activity. Genes Dev 20, 16–21 (2006). 10.1101/gad.1378506

37 Matheu, A. et al. Delayed ageing through damage protection by the Arf/p53 pathway. Nature 448, 375–379 (2007). 10.1038/nature05949

38 Donehower, L. A. et al. Mice deficient for p53 are developmentally normal but susceptible to spontaneous tumours. Nature 356, 215–221 (1992). 10.1038/356215a0

39 Jacks, T. et al. Tumor spectrum analysis in p53-mutant mice. Curr Biol 4, 1–7 (1994). 10.1016/s0960-9822(00)00002-6

40 Tanaka, H. et al. Nucleosome stability safeguards cell identity, stress resilience and healthy aging. bioRxiv (2025). 10.1101/2025.09.17.676776

41 Moreno, T. M. et al. Autophagy Flux Is Remodeled Sex- and Cell Type-Specifically During Human Aging, and Is Linked to Reduced Physical Function in Older Adults. Aging Cell 25, e70734 (2026). 10.1111/acel.70734

42 Sarkar, T. J. et al. Transient non-integrative expression of nuclear reprogramming factors promotes multifaceted amelioration of aging in human cells. Nat Commun 11, 1545 (2020). 10.1038/s41467-020-15174-3

43 Fischer, M. Census and evaluation of p53 target genes. Oncogene 36, 3943–3956 (2017). 10.1038/onc.2016.502

44 Chen, K. H., Boettiger, A. N., Moffitt, J. R., Wang, S. & Zhuang, X. RNA imaging. Spatially resolved, highly multiplexed RNA profiling in single cells. Science 348, aaa6090 (2015). 10.1126/science.aaa6090

45 Ran, F. A. et al. In vivo genome editing using Staphylococcus aureus Cas9. Nature 520, 186–191 (2015). 10.1038/nature14299

46 Sinha, S. et al. Aging disrupts hepatocyte zonation homeostasis in mice and humans. Hepatology 83, 1143–1157 (2026). 10.1097/HEP.0000000000001451

47 Whitfield, M. L. et al. Identification of genes periodically expressed in the human cell cycle and their expression in tumors. Mol Biol Cell 13, 1977–2000 (2002). 10.1091/mbc.02-02-0030

48 Subramanian, A. et al. Gene set enrichment analysis: a knowledge-based approach for interpreting genome-wide expression profiles. Proc Natl Acad Sci U S A 102, 15545–15550 (2005). 10.1073/pnas.0506580102

49 Herkel, J. et al. MHC class II-expressing hepatocytes function as antigen-presenting cells and activate specific CD4 T lymphocyutes. Hepatology 37, 1079–1085 (2003). 10.1053/jhep.2003.50191

50 He, S. et al. High-plex imaging of RNA and proteins at subcellular resolution in fixed tissue by spatial molecular imaging. Nat Biotechnol 40, 1794–1806 (2022). 10.1038/s41587-022-01483-z

51 Jin, S. et al. Inference and analysis of cell-cell communication using CellChat. Nat Commun 12, 1088 (2021). 10.1038/s41467-021-21246-9

52 Bernhagen, J. et al. MIF is a noncognate ligand of CXC chemokine receptors in inflammatory and atherogenic cell recruitment. Nat Med 13, 587–596 (2007). 10.1038/nm1567

53 Zhu, C. et al. The Tim-3 ligand galectin-9 negatively regulates T helper type 1 immunity. Nat Immunol 6, 1245–1252 (2005). 10.1038/ni1271

54 Fornes, O. et al. JASPAR 2020: update of the open-access database of transcription factor binding profiles. Nucleic Acids Res 48, D87–D92 (2020). 10.1093/nar/gkz1001

55 Whitehead, J. C. et al. A clinical frailty index in aging mice: comparisons with frailty index data in humans. J Gerontol A Biol Sci Med Sci 69, 621–632 (2014). 10.1093/gerona/glt136

56 Park, E. J. et al. Dietary and genetic obesity promote liver inflammation and tumorigenesis by enhancing IL-6 and TNF expression. Cell 140, 197–208 (2010). 10.1016/j.cell.2009.12.052

57 Havas, A. P. et al. Activated interferon signaling suppresses age-dependent liver cancer. bioRxiv (2024). 10.1101/2024.07.31.606057

58 Reyes, J. et al. Oncogenic and tumor-suppressive forces converge on a progenitor niche at the benign-to-malignant transition. Cell 189, 2875–2897 (2026). 10.1016/j.cell.2026.03.032

59 Dharel, N. et al. Potential contribution of tumor suppressor p53 in the host defense against hepatitis C virus. Hepatology 47, 1136–1149 (2008). 10.1002/hep.22176

60 Muñoz-Fontela, C. et al. Transcriptional role of p53 in interferon-mediated antiviral immunity. J Exp Med 205, 1929–1938 (2008). 10.1084/jem.20080383

61 Mackenzie, K. J. et al. cGAS surveillance of micronuclei links genome instability to innate immunity. Nature 548, 461–465 (2017). 10.1038/nature23449

62 Glück, S. et al. Innate immune sensing of cytosolic chromatin fragments through cGAS promotes senescence. Nat Cell Biol 19, 1061–1070 (2017). 10.1038/ncb3586

63 Dou, Z. et al. Cytoplasmic chromatin triggers inflammation in senescence and cancer. Nature 550, 402–406 (2017). 10.1038/nature24050

64 Harding, S. M. et al. Mitotic progression following DNA damage enables pattern recognition within micronuclei. Nature 548, 466–470 (2017). 10.1038/nature23470

65 Vizioli, M. G. et al. Mitochondria-to-nucleus retrograde signaling drives formation of cytoplasmic chromatin and inflammation in senescence. Genes Dev 34, 428–445 (2020). 10.1101/gad.331272.119

66 Komarova, E. A. et al. p53 is a suppressor of inflammatory response in mice. FASEB J 19, 1030–1032 (2005). 10.1096/fj.04-3213fje

67 Liu, G., Park, Y. J., Tsuruta, Y., Lorne, E. & Abraham, E. p53 Attenuates lipopolysaccharide-induced NF-kappaB activation and acute lung injury. J Immunol 182, 5063–5071 (2009). 10.4049/jimmunol.0803526

68 Tebaldi, T. et al. Whole-genome cartography of p53 response elements ranked on transactivation potential. BMC Genomics 16, 464 (2015). 10.1186/s12864-015-1643-9

69 Nguyen, T. T. et al. Revealing a human p53 universe. Nucleic Acids Res 46, 8153–8167 (2018). 10.1093/nar/gky720

70 Yahagi, N. et al. p53 activation in adipocytes of obese mice. J Biol Chem 278, 25395–25400 (2003). 10.1074/jbc.M302364200

71 Jiang, P. et al. p53 regulates biosynthesis through direct inactivation of glucose-6-phosphate dehydrogenase. Nat Cell Biol 13, 310–316 (2011). 10.1038/ncb2172

72 Assaily, W. et al. ROS-mediated p53 induction of Lpin1 regulates fatty acid oxidation in response to nutritional stress. Mol Cell 44, 491–501 (2011). 10.1016/j.molcel.2011.08.038

73 Prokesch, A. et al. Liver p53 is stabilized upon starvation and required for amino acid catabolism and gluconeogenesis. FASEB J 31, 732–742 (2017). 10.1096/fj.201600845R

74 Porteiro, B. et al. Pharmacological stimulation of p53 with low-dose doxorubicin ameliorates diet-induced nonalcoholic steatosis and steatohepatitis. Mol Metab 8, 132–143 (2018). 10.1016/j.molmet.2017.12.005

75 Porteiro, B. et al. Hepatic p63 regulates steatosis via IKKβ/ER stress. Nat Commun 8, 15111 (2017). 10.1038/ncomms15111

76 Tomita, K. et al. p53/p66Shc-mediated signaling contributes to the progression of non-alcoholic steatohepatitis in humans and mice. J Hepatol 57, 837–843 (2012). 10.1016/j.jhep.2012.05.013

77 Derdak, Z. et al. Inhibition of p53 attenuates steatosis and liver injury in a mouse model of non-alcoholic fatty liver disease. J Hepatol 58, 785–791 (2013). 10.1016/j.jhep.2012.11.042

78 Tschaharganeh, D. F. et al. p53-dependent Nestin regulation links tumor suppression to cellular plasticity in liver cancer. Cell 158, 579–592 (2014). 10.1016/j.cell.2014.05.051

79 Morris, J. P. et al. α-Ketoglutarate links p53 to cell fate during tumour suppression. Nature 573, 595–599 (2019). 10.1038/s41586-019-1577-5

80 Jeay, S. et al. Dose and Schedule Determine Distinct Molecular Mechanisms Underlying the Efficacy of the p53-MDM2 Inhibitor HDM201. Cancer Res 78, 6257–6267 (2018). 10.1158/0008-5472.CAN-18-0338

81 Lane, N. E. et al. A phase 2, randomized, double-blind, placebo-controlled study of senolytic molecule UBX0101 in the treatment of painful knee osteoarthritis. Osteoarthritis Cartilage 29, S52–S53 (2021). 10.1016/j.joca.2021.02.077

82 Charni-Natan, M. & Goldstein, I. Protocol for Primary Mouse Hepatocyte Isolation. STAR Protoc 1, 100086 (2020). 10.1016/j.xpro.2020.100086

83 Conant, D. et al. Inference of CRISPR Edits from Sanger Trace Data. CRISPR J 5, 123–130 (2022). 10.1089/crispr.2021.0113

84 Goode, A., Gilbert, B., Harkes, J., Jukic, D. & Satyanarayanan, M. OpenSlide: A vendor-neutral software foundation for digital pathology. J Pathol Inform 4, 27 (2013). 10.4103/2153-3539.119005

85 Martin, M. Cutadapt removes adapter sequences from high-throughput sequencing reads. *EMBnet.journal* 17, 10–12 (2011). 10.14806/ej.17.1.200

86 Dobin, A. et al. STAR: ultrafast universal RNA-seq aligner. Bioinformatics 29, 15–21 (2013). 10.1093/bioinformatics/bts635

87 Heinz, S. et al. Simple combinations of lineage-determining transcription factors prime cis-regulatory elements required for macrophage and B cell identities. Mol Cell 38, 576–589 (2010). 10.1016/j.molcel.2010.05.004

88 Love, M. I., Huber, W. & Anders, S. Moderated estimation of fold change and dispersion for RNA-seq data with DESeq2. Genome Biol 15, 550 (2014). 10.1186/s13059-014-0550-8

89 Wu, T. et al. clusterProfiler 4.0: A universal enrichment tool for interpreting omics data. Innovation (Camb) 2, 100141 (2021). 10.1016/j.xinn.2021.100141

90 Korotkevich, G. et al. Fast gene set enrichment analysis. bioRxiv (2021). 10.1101/060012

91 Hunter, J. D. Matplotlib: A 2D graphics environment. Computing in Science & Engineering 9, 90–95 (2007). 10.1109/MCSE.2007.55

92 Virtanen, P. et al. SciPy 1.0: fundamental algorithms for scientific computing in Python. Nat Methods 17, 261–272 (2020). 10.1038/s41592-019-0686-2

93 Fleming, S. J. et al. Unsupervised removal of systematic background noise from droplet-based single-cell experiments using CellBender. Nat Methods 20, 1323–1335 (2023). 10.1038/s41592-023-01943-7

94 McGinnis, C. S., Murrow, L. M. & Gartner, Z. J. DoubletFinder: Doublet Detection in Single-Cell RNA Sequencing Data Using Artificial Nearest Neighbors. Cell Syst 8, 329–337 e324 (2019). 10.1016/j.cels.2019.03.003

95 Hao, Y. et al. Dictionary learning for integrative, multimodal and scalable single-cell analysis. Nat Biotechnol 42, 293–304 (2024). 10.1038/s41587-023-01767-y

96 Stuart, T., Srivastava, A., Madad, S., Lareau, C. A. & Satija, R. Single-cell chromatin state analysis with Signac. Nat Methods 18, 1333–1341 (2021). 10.1038/s41592-021-01282-5

97 Korsunsky, I. et al. Fast, sensitive and accurate integration of single-cell data with Harmony. Nat Methods 16, 1289–1296 (2019). 10.1038/s41592-019-0619-0

98 Stringer, C., Wang, T., Michaelos, M. & Pachitariu, M. Cellpose: a generalist algorithm for cellular segmentation. Nat Methods 18, 100–106 (2021). 10.1038/s41592-020-01018-x

99 Hafemeister, C. & Satija, R. Normalization and variance stabilization of single-cell RNA-seq data using regularized negative binomial regression. Genome Biol 20, 296 (2019). 10.1186/s13059-019-1874-1

100 Patil, A. & Patil, A. CellKb Immune: a manually curated database of mammalian hematopoietic marker gene sets for rapid cell type identification. bioRxiv (2020). 10.1101/2020.12.01.389890

101 Bankhead, P. et al. QuPath: Open source software for digital pathology image analysis. Sci Rep 7, 16878 (2017). 10.1038/s41598-017-17204-5

102 Wolock, S. L., Lopez, R. & Klein, A. M. Scrublet: Computational Identification of Cell Doublets in Single-Cell Transcriptomic Data. Cell Syst 8, 281–291 e289 (2019). 10.1016/j.cels.2018.11.005

103 Hao, Y. et al. Integrated analysis of multimodal single-cell data. Cell 184, 3573–3587 (2021). 10.1016/j.cell.2021.04.048

104 McInnes, L., Healy, J. & Melville, J. UMAP: uniform manifold approximation and projection for dimension reduction. arXiv (2018). 10.48550/arXiv.1802.03426

105 Jin, S., Plikus, M. V. & Nie, Q. CellChat for systematic analysis of cell-cell communication from single-cell transcriptomics. Nat Protoc 20, 180–219 (2025). 10.1038/s41596-024-01045-4

